# Three-dimensional quantitative micro-elastography reveals alterations in spatial elasticity patterns of follicles and corpora lutea in murine ovaries during ageing

**DOI:** 10.1101/2024.10.08.617177

**Authors:** Anna Jaeschke, Matt S. Hepburn, Alireza Mowla, Brendan F. Kennedy, Chii Jou Chan

## Abstract

Fibrosis and tissue stiffening are hallmarks of ovarian ageing, linked to a decrease in fertility. However, the lack of three-dimensional (3D) characterization of ovary elasticity limits our understanding of localized elasticity patterns and their connection to the tissue composition. Here, we developed an integrated approach to link ovarian tissue elasticity, volume, and cell-matrix composition using quantitative micro-elastography (QME), a label-free, non-invasive approach to study the 3D microscale elasticity in conjunction with immunofluorescence microscopy. QME revealed distinct spatial elasticity patterns in ovarian compartments, namely follicles and corpora lutea (CLs), and local elasticity alterations in different age cohorts. For example, CL elasticity significantly increased during ovarian ageing while follicle elasticity changed minimally. CLs showed size-dependent elasticity changes, while follicles exhibited distinct spatial variations in elasticity correlated with the emergence of theca cell layers during follicle development. These findings have the potential to guide the development of novel diagnostic tools and identify therapeutic targets, improving women’s reproductive health and longevity.

**Graphical Abstract:** 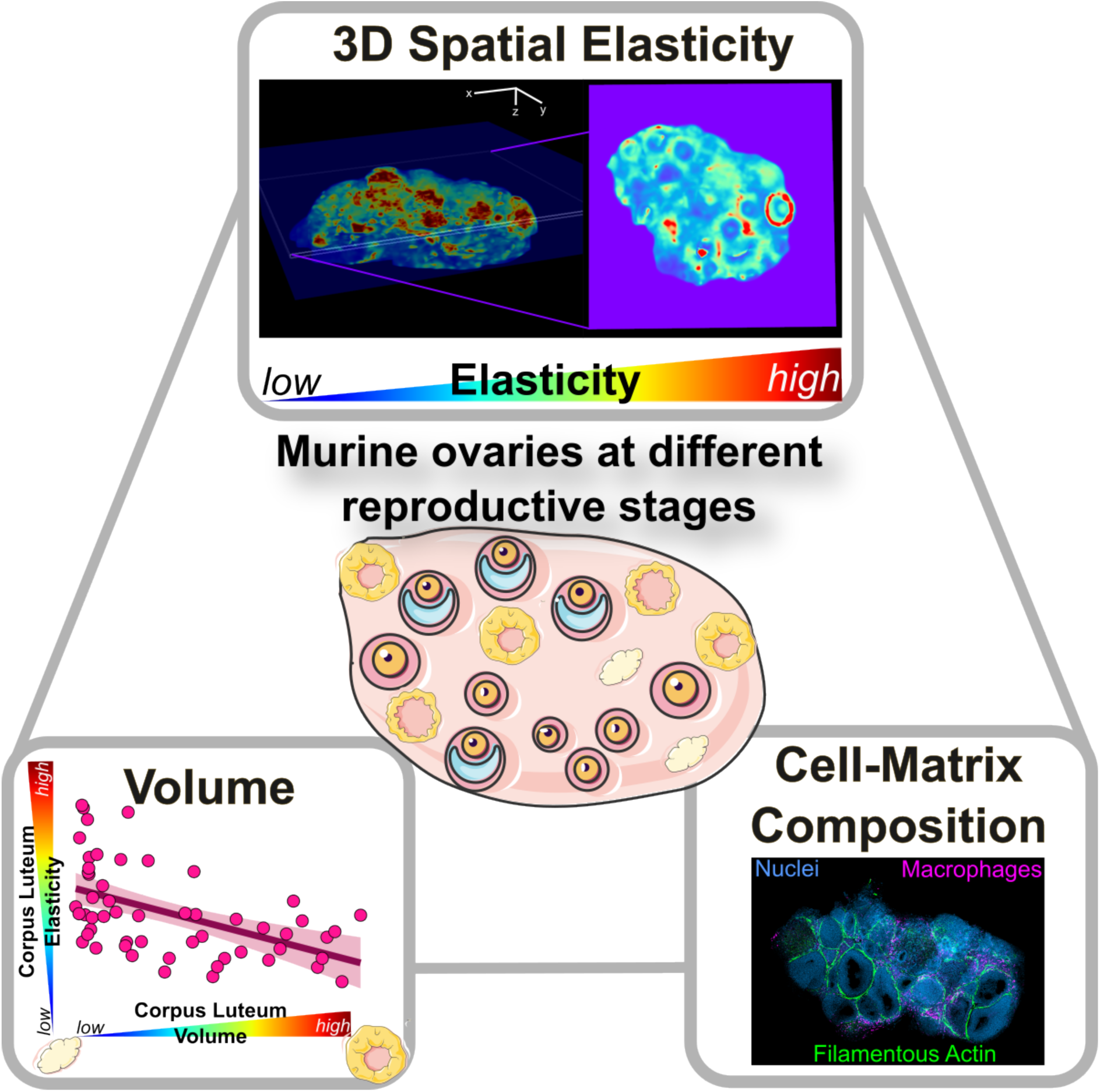

## INTRODUCTION

The decline in female fertility with age is well-established and is attributed to a reduction in follicle number and quality, posing challenges to the effectiveness of *in vitro* fertilization. Within the ovary, follicles are embedded in an extracellular matrix (ECM) and surrounded by a myriad of highly specific cell types. Each component of the ovarian tissue architecture is crucial for tissue homeostasis and function^1^. The mechanical microenvironment, alongside hormonal regulation, contributes significantly to regulating follicle development^2,3^. For example, rigid microenvironments support early-stage follicle survival and growth, while softer matrices promote late-stage antrum formation *in vitro*^4–7^. Changes in the composition and mechanical properties of the tissue are hallmarks of ovarian ageing. Increased collagen and decreased hyaluronan content have been associated with elevated tissue elasticity^8–11^ contributing to mis-ovulation and decreased fertility^12–14^. Targeted removal of fibrotic ECM has been shown to maintain fertility in aged mice^13,15^, emphasizing the significance of the mechanical microenvironment for ovarian function.

The ovarian compartments, including follicles, corpora lutea (CLs), and stroma, have distinct functional roles within the ovary. While there have been several studies on ovarian elasticity^8,11,16^, how follicle and CL elasticity may independently contribute to ovarian stiffening during ageing remains understudied. CLs, originating from the remnants of ovarian follicles post-ovulation, undergo rapid cellular remodeling, growth, and eventual regression linked to prominent changes in ECM composition^17–21^. CLs at various stages of development may exhibit distinct mechanical properties that contribute to increased tissue heterogeneity in the ovary, although this hypothesis has yet to be tested. Furthermore, recent studies have shown that oocytes are sensitive to mechanical stress^22^. Therefore, studying intra-follicle elasticity patterns during development is essential to understanding how the mechanical environment impacts oocyte maturation and quality at different reproductive ages.

Various techniques have been used to study the mechanical phenotype of ovarian tissues on the microscale. Multi-modal indentation-based approaches have revealed increasing organ elasticity upon ageing^8,11^. Measurements across the organ have uncovered rigidity gradients between the outer layer (cortex) and inner part (medulla) as well as intra-follicular mechanical heterogeneities^23,24^. Using Brillouin microscopy, Chan *et al*. revealed the presence of a mechanically stiff follicle ‘shell’ that emerges during follicle development^25^. Despite providing important insights into ovarian function, existing techniques have several notable limitations. For example, sample preparation often involves bisecting or sectioning the organ, significantly altering its mechanical environment. Furthermore, low scan acquisition speed and limited effective penetration depth restrict the ability to characterize the tissue mechanical phenotype in three dimensions (3D) and link elasticity patterns to the cellular phenotypes and matrix composition.

In this study, we applied quantitative micro-elastography (QME)^26^, a variant of compression-based optical coherence elastography, to characterize the 3D microscale elasticity of intact ovaries. We developed an integrated method to generate volumetric images of ovarian microscale elasticity co-registered with immunofluorescence microscopy, linking tissue elasticity, volume, and cell-matrix composition. Using this approach, we characterized changes in elasticity, volume, and cell-matrix composition of murine ovaries across three different age cohorts, each corresponding to a different reproductive stage. Mice at 3 weeks of age represent the pre-pubertal stage before the onset of cyclicity (first ovulation)^27^. At the age of 9 weeks, mice were in the reproductively active phase. The 12-month cohort exhibited longer cycles and hormonal fluctuations linked to the onset of ovarian senescence and follicle depletion, which are similar to the changes observed in humans at perimenopause^28^. Integrating QME with light microscopy imaging techniques, we identified distinct 3D spatial elasticity patterns in the ovaries, corresponding to follicles and CLs. Our findings reveal a significant increase in CL elasticity with age, while follicle elasticity remains relatively stable, suggesting that CLs are key contributors to age-related changes in ovarian tissue elasticity. We further established that CL elasticity correlates closely with their volume, cellular phenotypes, and matrix composition. Moreover, we uncovered emerging spatial elasticity patterns within follicles that correlate with their developmental stages. We anticipate that extending this work to studying ovarian disease and infertility models could facilitate the identification of novel biomarkers and therapeutics for fibrotic diseases.

## RESULTS

### Volumetric imaging of microscale elasticity in murine ovaries of different reproductive age cohorts

Using QME, we assessed the 3D spatial-resolved local elasticity of murine ovaries *ex vivo*. Figure 1 presents optical coherence tomography (OCT) and OCT/QME overlay images of one representative ovary for each age group. OCT revealed internal ovarian structures such as follicles, oocytes, and CLs (Figures 1A,B). The tissue was highly scattering, and the OCT depth penetration varied between samples due to differences in ovary morphologies and material properties. However, in all tissues, we achieved a high OCT signal-to-noise ratio (SNR) of ≥ 10 decibels (dB) at depths of at least 500 μm (Figure 1; SVideos 1-6). For elasticity quantification, we defined an ‘imaging volume’ as the ovary volume measured to a constant depth of 500 μm below the ovary’s top surface to ensure consistent data quality.

**Figure 1.**
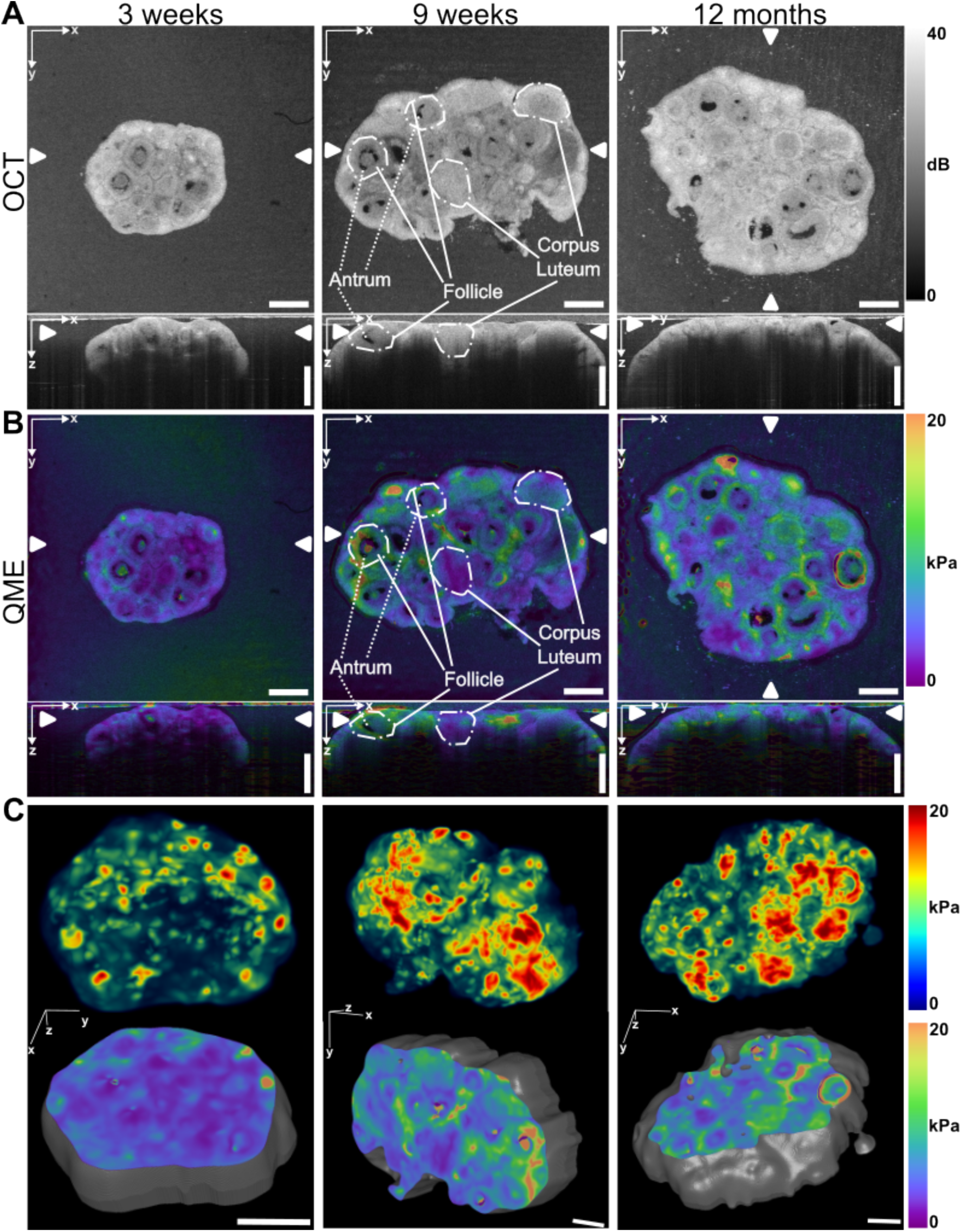
Volumetric imaging of microscale elasticity in murine ovaries at different reproductive stages: A) OCT images of one representative murine ovary at 3 weeks, 9 weeks, and 12 months. B) Corresponding OCT/QME overlay images. The locations of the cross-sections in the corresponding orthogonal planes are indicated by white arrowheads. C) Views of a 3D rendering (top row) and z-slice (bottom row) showing internal cross-sectional detail and outline of the organ (gray) of QME images for each ovary. Animations of the 3D reconstructions are presented in SVideo 4-6. Scale bars: 500 µm.

We examined the elasticity of ovaries from mice in three age groups: 3 weeks, 9 weeks, and 12 months, representing pre-puberty, reproductive activity, and decline in fertility, respectively. The QME images revealed non-uniform elasticity across the entire ovaries, which is partly associated with follicles of various sizes and stages (Figures 1B,C). As each voxel represents an elasticity measurement, the QME quantification resulted in several million data points per sample. Pooling data from all five ovaries within each age group revealed a right-skewed distribution (Figures 2A, S1A). We described the distributions and compared elasticity in different groups using peak elasticity (mode of the elasticity distribution) and median elasticity measurements for each ovary across the age cohorts (Figures 2B, S1B).

**Figure 2.**
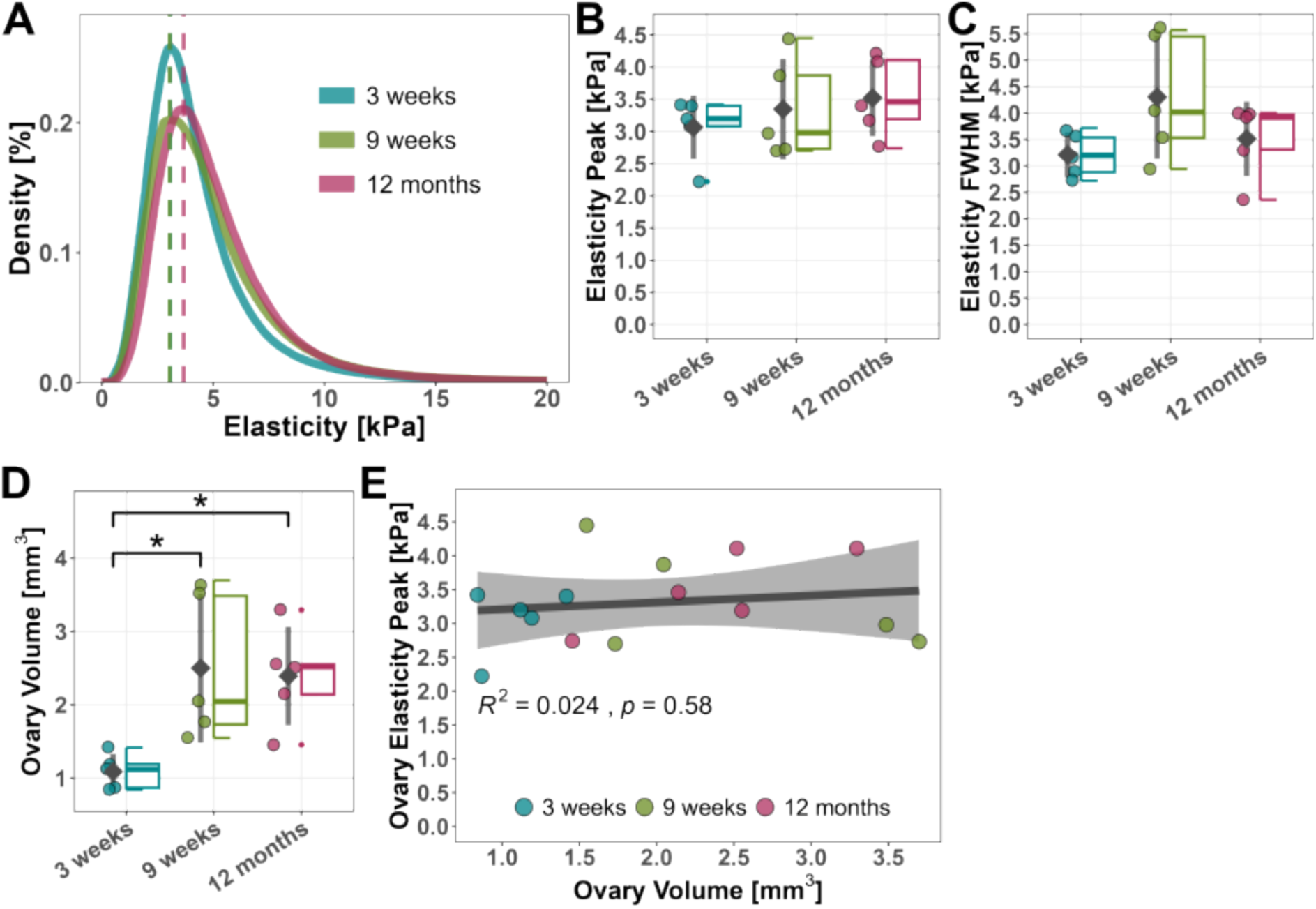
Total murine ovary elasticity at different reproductive stages: A) Distribution of elasticity measurements of all ovaries pooled per age group; dashed lines indicate peak elasticity for each age group. B) Elasticity peak and C) FWHM for each ovary across the different age groups. D) Volume measurements for each ovary across the different age groups. E) Correlation between peak organ elasticity and organ volume. B-E) n = 5 ovaries per age group. B-D) Statistical significance between mean values determined by a one-way ANOVA D) followed by a Tukey’s post-hoc test (*p < 0.05). B,C) No significant differences were found between the mean values of the age groups.

We observed a slight increase in peak elasticity from 3.06 ± 0.49 kPa for ovaries at 3 weeks to 3.35 ± 0.78 kPa at 9 weeks and 3.52 ± 0.6 kPa at 12 months (mean ± standard deviation of peak elasticities from 5 ovaries per age group; Figure 2B). Similarly, the median elasticity changed from 3.77 ± 0.68 kPa in 3-week-old ovaries to 4.45 ± 0.81 kPa (9 weeks) and 4.40 ± 0.69 kPa (12 months; Figure S1B). Furthermore, we observed a broader elasticity distribution for the 9-week and 12-month ovaries (Figures 2A, S1A). Such variance was observed not only between age cohorts but also between samples within the same age group (Figure S1D). The elasticity full width at half maximum (FWHM) changed from 3.21 ± 0.42 kPa at 3 weeks to 4.30 ± 1.17 kPa at 9 weeks and 3.51 ± 0.704 kPa at 12 months, indicating a slightly increased tissue elasticity heterogeneity in older ovaries (Figure 2C). The trend in the intra-group heterogeneity was further supported by an increase in elasticity interquartile range (IQR) values from 3 weeks to 9 weeks and 12 months (Figure S1C). We explored the relationship between organ volume and elasticity and found a significant increase in ovarian volume from 3 weeks to 9 weeks and 12 months (Figure 2D), capturing the organ’s growth between the ages of 3 weeks and 9 weeks. However, interestingly, we found no significant correlation between the total ovarian elasticity and their ‘imaging volume’ (Figures 2E, S1E), suggesting minimal changes in total ovary elasticity with age.

Notably, despite only subtle changes in the total ovary elasticity between age groups, the QME images showed distinct, local variations in elasticity in the different ovarian structures (Figure 1B). This suggests that computing an average elasticity value for an entire organ does not capture significant spatial variations in elasticity that influence ovarian function. QME allowed us to further study the spatial variations in tissue elasticity, a key advantage over existing bulk elasticity measurement techniques. Combining QME with light microscopy, we could identify functional compartments in the ovary. We co-registered the regions of interest (ROIs) from the light microscopy to the QME data set to extract elasticity and volume, focusing on CLs and follicles, the main tissue components. Using this approach, we found that the total number of follicles and CLs was similar between ovaries in different age groups (Figures S1F,G (left panels)). Defining CL and follicle volume ratios, by dividing the total CL or follicle volume by the ‘imaging volume’ of ovaries, we found that the follicle volume fraction was considerably smaller in 9-week and 12-month ovaries compared to 3-week-old ovaries (Figure S1G (middle and right panel)). However, there were no significant differences in CL or follicle volume fractions between 9-week and 12-month ovaries (Figures S1F,G (middle and right panel)). We did not observe a correlation between total ovary elasticity and the volume fraction occupied by CLs or follicles (Figures S1F,G (middle and right panel)), suggesting that additional factors, such as the intrinsic elasticity of CLs or follicles, may contribute to total ovary elasticity.

### CL elasticity is a key contributor to elasticity in aged ovaries

We examined CL elasticity in the 9-week and 12-month ovaries (Figures 3A,B). The 3-week-old mice were in the pre-puberty phase, not undergoing estrous cycles, and therefore, no CLs were formed in the ovaries. The CL elasticity followed a similar right-skewed distribution as the whole organ elasticity (Figure 3B). We observed a notable increase in the pooled elasticity of all CLs between ovaries at 9 weeks and 12 months (Figures 3C, S2A). Peak elasticity increased from 4.19 ± 1.59 kPa in 9-week-old CLs to 5.22 ± 2.02 kPa in 12-month-old CLs (Figure 3D). Similarly, median elasticity changed from 4.83 ± 1.47 kPa to 5.88 ± 2.09 kPa (Figure S2B).

**Figure 3.**
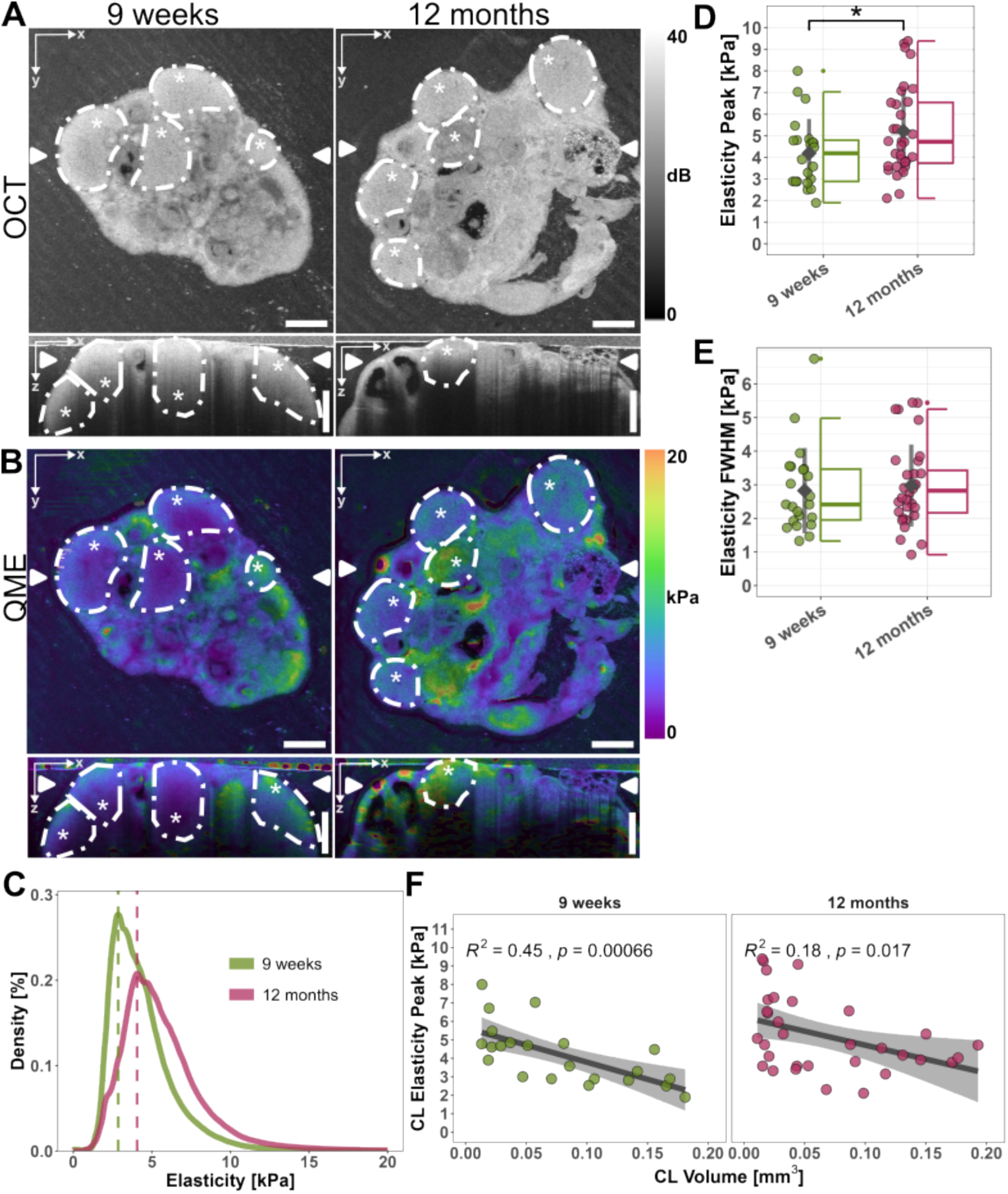
Increased elasticity of CLs in aged ovaries: A) Representative OCT and B) QME images of CLs in 9-week and 12-month-old ovaries. Dashed lines and asterisks indicate the segmented CLs. Scale bars: 500 µm. C) Distribution of elasticity measurements of all CLs pooled per age group; lines indicate peak elasticity for each age group. D) Elasticity peak and E) FWHM for each CL across the different age groups. E) Correlation between CL peak elasticity and CL volume across the different age groups. B-E) n = 22 CLs from 4 ovaries (9 weeks) and 32 CLs from 5 ovaries (12 months). D,E) Statistical significance between mean values determined by a Student’s t-test (*p < 0.05). E) No significant differences were found between the mean values of the age groups.

Focusing on individual CL elasticity, we observed considerable variance in both age groups but no significant changes in heterogeneity between the age groups (Figure 3E, S2C). Using the 3D capability of QME, we found that CL elasticity decreased with increasing size in the 9-week ovaries (Figures 3F and S2D, left panels). At the same time, a similar trend was observed in the 12-month ovaries with greater variability (Figure 3F and S2D, right panels). Notably, the 12-month ovaries were populated with a higher number of small and stiff CLs, contributing to the higher overall CL elasticity in the older ovaries.

We used immunofluorescence (IF) to investigate the cell-molecular composition of CLs in 9-week-old and 12-month-old ovaries as a potential cause of elasticity alterations. We found no difference in cell number per area between young and aged CLs (Figure 5A, left panel). Moreover, there was no correlation between cell number per area and elasticity, indicating that elasticity changes are unrelated to cell packing (Figure 5A, right panel). A reduced number of alpha-smooth muscle actin (α-SMA) expressing cells suggested fewer cells with a contractile phenotype in aged CLs (Figures 4A, 5B, S3A). Similarly, reduced filamentous actin (F-actin) content indicated decreased stress fiber formation within aged CLs (Figures 4A,B, 5C).

**Figure 4.**
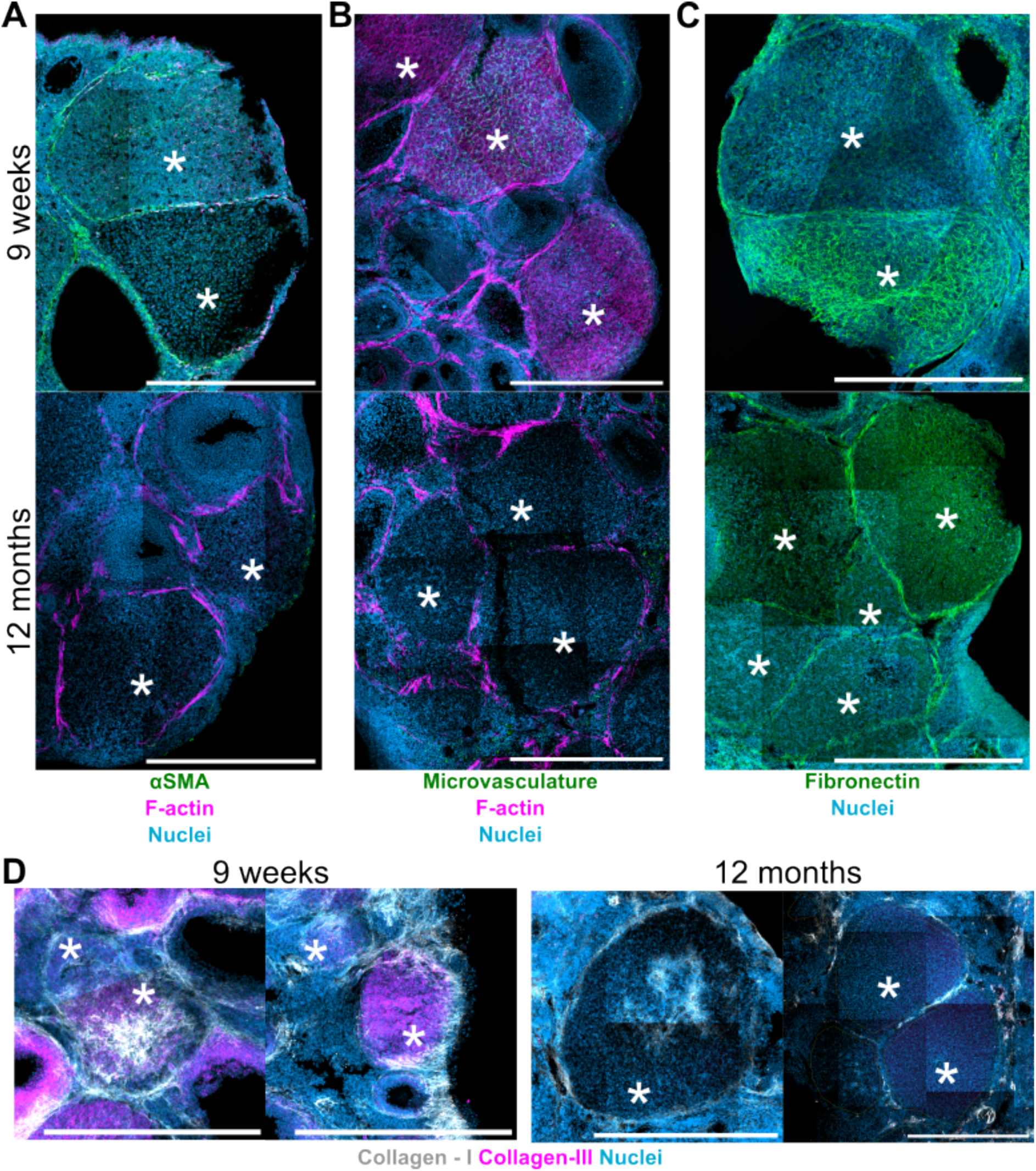
Cellular phenotypes and ECM composition of CLs in 9-week and 12-month-old ovaries: Representative IF images of A) αSMA (green), F-actin (magenta), B) microvasculature (CD31; green), F-actin (magenta), C) Fibronectin (green), D) collagen-I (grey), and collagen-III (magenta). A-D) Nuclei in cyan. Asterisks indicate CLs. Scale bars: 500 µm.

**Figure 5.**
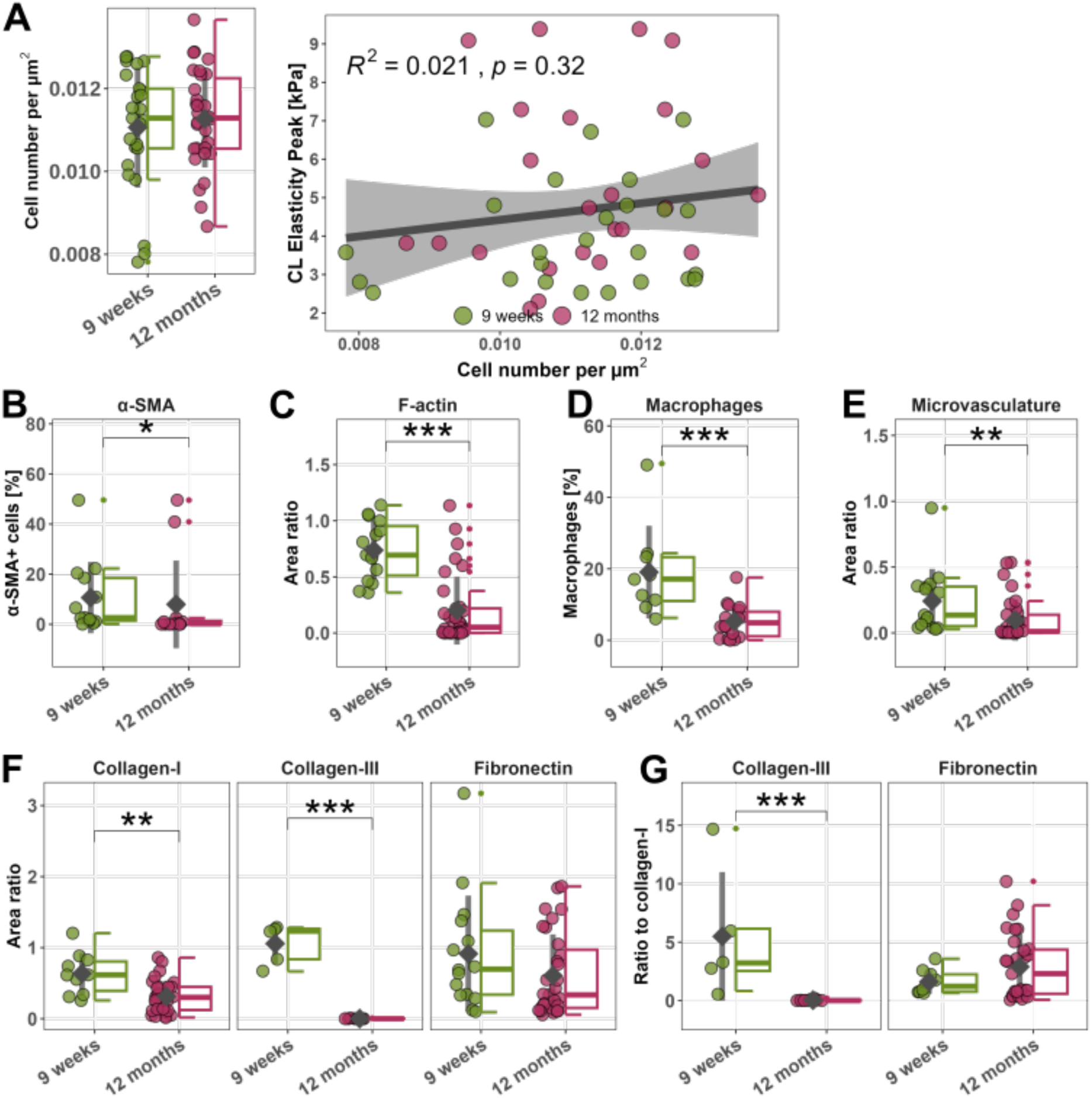
Changes in cell and molecular composition in young and aged CLs: A) Cell number per µm^2^ in young and aged CL (left panel) and correlation with peak elasticity (right panel), n = 25 CLs from 3 ovaries (9 weeks) and 24 CLs from 4 ovaries (12 months). B) Percentage of α-SMA expressing cells in young and aged CLs, n = 13 CLs from 3 ovaries (9 weeks) and 12 CLs from 2 ovaries (12 months); C) F-actin positive area in CLs, n = 15 CLs from 3 ovaries (9 weeks) and 33 CLs from 4 ovaries (12 months); D) Percentage of macrophages (F4/80 expressing cells) in young and aged CLs; n = 9 CLs from 2 ovaries (9 weeks) and 18 CLs from 3 ovaries (12 months); E) Microvasculature (CD31) positive area in young and aged CLs, n = 15 CLs from 3 ovaries (9 weeks) and 33 CLs from 4 ovaries (12 months); F) Collagen-I, n = 10 CLs from 2 ovaries (9 weeks) and 31 CLs from 4 ovaries (12 months); collagen-III, n = 5 CLs from 1 ovaries (9 weeks) and 13 CLs from 2 ovaries (12 months); and fibronectin, n = 15 CLs from 3 ovaries (9 weeks) and 31 CLs from 4 ovaries (12 months); positive area, in young and aged CLs. G) Collagen-III, n = 5 CLs from 1 ovary (9 weeks) and 13 CLs from 2 ovaries (12 months); and fibronectin, n = 9 CLs from 2 ovaries (9 weeks) and 31 CLs from 4 ovaries (12 months); area ratio normalized to collagen-I area ratio of each CL. C,E,F,G) area ratios of markers normalized by the area ratio occupied by nuclei. A,C-G) Statistical significance between distributions determined by a Mann-Whitney-U-test (*p < 0.05, ** p < 0.01, ***p < 0.001).

Interestingly, while the F-actin expression was reduced inside the aged CLs, an F-actin-rich ring surrounding the CLs remained (Figures 4 A,B). We observed increased stress fiber content with increasing volume in young CLs but not aged ones (Figure S3B, right panel) and a decrease in the total macrophage count (F4/80 expressing cells) within the CLs in young versus old mice (Figure 5D). Interestingly, the macrophage count was greatly reduced in larger CLs in the 9-week cohort (Figure S3C, right panel). Moreover, the microvessel area decreased in aged CLs (Figures 4B, 5E) but showed no correlation with volume or elasticity (Figure S3D).

Aged CLs exhibited lowered collagen-I and collagen-III content (Figures 4D, 5F), while fibronectin levels remained similar between CLs of the two age cohorts. (Figures 4C, 5F). In 9-week-old ovaries, CLs of higher elasticity showed a trend of decreased collagen-III and fibronectin content (Figures S4B,C), a pattern not observed for collagen-I (Figure S4A). Besides the expression of individual ECM constituents, the proportion of matrix components is relevant for tissue function. The lowered collagen-III to collagen-I ratio (Figure 5G) in aged CLs was consistent with the low expression of collagen-III in aged CLs, emphasizing that collagen-III decreased more significantly than collagen-I during ageing. The fibronectin to collagen-I ratio was slightly increased in aged CLs (Figure 5G), indicating that fibronectin remained relatively stable between young and aged CLs while collagen-I expression decreased with ageing.

The heterogeneity in cellular and ECM composition was notable not only between the two age cohorts but also within ovaries of the same age. In the same ovary, CLs in proximity showed significant variations in the cellular and matrix composition. For example, in Figure 4A, in the 9-week ovary, microvessels were present in one CL but not the adjacent one. Similar patterns were found for fibronectin (Figure 4C). Analyzing the ECM, we observed CLs that were rich in both collagen-I and collagen-III. In contrast, others were only rich in collagen-III, lacking collagen-I (Figure 4D, compare left and right images of the 9-week panel). In 12-month-old ovaries, all CLs lacked collagen-III, but some were rich in collagen-I, and others were not (Figure 4D, compare left and right images of the 12-week panel).

Taken together, these results suggest that alterations in cellular phenotypes and ECM composition may contribute to changes in tissue elasticity of aged CLs.

### Spatial mechanical patterns in follicles at different developmental stages

We next examined if there are changes to follicle elasticity during ageing. We observed only a minimal increase in follicle peak elasticity between different age groups, measuring 3.62 ± 0.90 kPa, 3.93 ± 0.91 kPa, and 4.09 ± 0.90 kPa at 3 weeks, 9 weeks, and 12 months, respectively, with similar elasticity distributions (Figure 6C). It is well reported that follicle development is accompanied by significant alterations in the cellular and molecular composition alongside an increase in size^29–31^. Quantitative observation from the QME images shows that follicles exhibit non-uniform elasticity. Specifically, we observed a ring of increased elasticity around some follicles (Figures 6A,B, middle panel, purple rectangle) but not in others (Figures 6A,B, middle panel, orange rectangle). We analyzed IF-stained ovaries and found that the high elasticity ring coincided with F-actin and α-SMA rich regions (Figures 6A,B, right panel). To investigate the spatial patterns of elasticity in more detail, we segmented the QME data using manually drawn concentric circles to separate the follicle elasticity measurements into ‘inner’ and ‘outer’ areas and the oocyte (Figure 6D). Frequently, the oocyte displayed high elasticity (Figure 6A, mid panel, purple rectangle; also Figure 1A). In late-stage follicles, the oocyte can become detached from the surrounding tissue due to the presence of follicular fluid, resulting in ineffective stress transmission from the actuator during elastography and minimal deformation. This invalidates the mechanical model used in QME image formation, as the artificially low strain in the oocyte from the ineffective stress transmission can lead to artificially high elasticity values representing an imaging artifact. Consequently, elasticity measurements in the oocyte area were excluded to avoid confounding the subsequent analyses.

**Figure 6.**
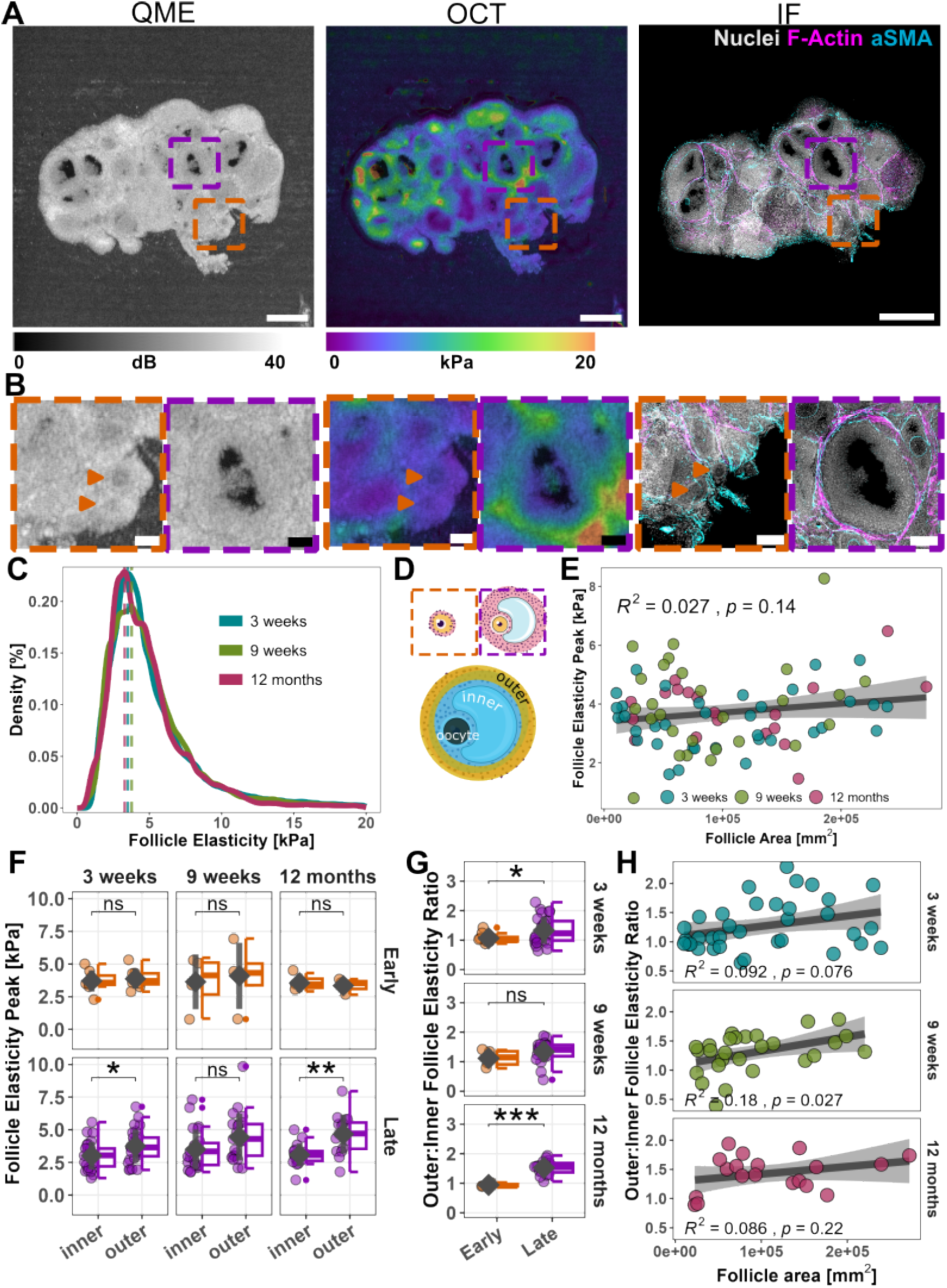
Compartments with varying elasticity emerge during follicle development: A) Representative OCT (left panel), QME (middle panel), IF (right panel) images of follicles in a 9-week-old ovary. Scale bar: 500 µm. B) Magnified regions of early (orange border) and late (purple border) stage follicles. Orange arrowheads indicate the location of two early-stage follicles. Scale bars: 100 µm. C) Distribution of elasticity measurements of all follicles pooled per age group; lines indicate elasticity peak for each age group; n=5 ovaries per age group. D) Schematic of early and late-stage follicles and segmentation approach of inner, outer, and oocyte regions. Illustrations adapted from Servier Medical Art under CC BY 4.0. E) Correlation between follicle elasticity peak and follicle area across the different age groups. F) Follicle elasticity peaks in the inner and outer segments of each age group. G) Ratio between outer and inner elasticity peak per follicle for early and late-stage follicles across the different age groups. H) Correlation between outer to inner elasticity peak ratio and follicle area in each age group. E-H) n= 16 early follicles, 54 late follicles from 5 ovaries (3 weeks), 8 early follicles, 46 late follicles from 4 ovaries (9 weeks), 6 early follicles, 32 late follicles from 5 ovaries (12 months).

We found no correlation between total follicle elasticity (excluding the oocyte) and follicle area (Figure 6E). To investigate follicles at different developmental stages, we broadly classified them into early-stage, small follicles and late-stage, larger follicles based on their area (Figure 6D) and the presence of a visible antrum in the OCT images. In early-stage follicles, the elasticity was similar between the inner and outer regions. In contrast, later-stage follicles showed a significant increase in the elasticity of the outer ring across all three age cohorts (Figure 6F). The elasticity ratio between the outer and inner segment elasticity was quantified for each follicle to investigate the change per follicle. The elasticity of the outer ring increased in late-stage follicles to about 2–2.5 times that of the inner elasticity (Figure 6G). Additionally, we found that the elasticity ratio of the outer to inner segments increased with follicle area in ovaries from 3-week and 9-week-old mice (Figure 6H), suggesting that the follicle shells become stiffer during development.

## DISCUSSION

In this study, we combined QME with 3D morphometric analysis and co-registered IF staining to track alterations in spatial elasticity patterns during ageing, linking them to the present cell-molecular phenotypes. While ultrasound elastography and magnetic resonance elastography can provide 3D tissue characterization *in vivo*, they are typically limited to spatial resolution greater than 1 mm^32,33^. Recently, ultrasound elastography has been used to probe human ovarian elasticity *in vivo*, reporting elasticity values of the same order of magnitude as our study^34^. However, with a spatial resolution of 35 µm, our QME setup allows us to map ovarian elasticity distributions in much finer detail. This higher resolution enabled us to examine microscale spatial elasticity patterns on length scales similar to those of cells, providing a deeper understanding of the mechanical properties of specific functional entities within the tissue. In contrast to surface indentation methods such as atomic force microscopy, QME is non-invasive and does not require tissue sectioning, preserving the native mechanical state of ovarian components. In recent years, Brillouin microscopy^35^ has emerged as a tool to probe tissue elasticity with high spatial resolution (0.5 – 1 µm) but has limited penetration depth. QME measures Young’s modulus directly and overcomes several limitations of other methods by avoiding phototoxicity in living samples and enabling rapid acquisition of volumetric elasticity images for the entire ovary within minutes. This makes QME a practical and efficient tool to characterize ovarian tissue in 3D with fields of view that extend across the entire organ with microscale spatial resolution.

Unlike indentation-based studies, we did not observe a significant increase in ovary elasticity with ageing^8^ or the emergence of an elasticity gradient between the cortex and the medullary region^23,24^. Instead, with QME, we observed non-uniform regions of elasticity across the organ linked to functional components. In line with previous studies^36,37^, we found a decrease in follicle numbers between reproductively active and aged mice, indicating a depletion in the follicle reserve with ageing. Interestingly, we saw increased ovary volume and overall elasticity variance between pre-puberty and older mice. Higher variations in elasticity have also been observed in skeletal muscle tissue with aging^38^. However, the lack of a prominent change in ovary volume and elasticity between 9-week and 12-month mice suggests that the presence of CLs significantly influences overall volume and tissue heterogeneity rather than ageing alone.

We propose that CLs are key contributors to ovarian elasticity, as they showed a pronounced increase in elasticity with aging compared to the total organ elasticity and follicles. Physiologically, the CL is an endocrine structure that produces hormones during pregnancy. Every time a follicle is ovulated, a CL is formed, which regresses either at the later stages of pregnancy or if pregnancy does not occur. Previous studies reported decreased CL numbers in aged ovaries^12,37^, which is in contrast to our study and can be caused by differences in mice strains and age cohorts. In multi-ovulatory species, CL volume and number are affected by factors such as mouse age, the number of previous litters, and the number of fetuses^39,40^. The high variance in CL number we observed between organs in the 12-month cohort reflects the dynamic changes during onset of ovarian senescence and decline of ovarian function.

The CL undergoes regression if pregnancy does not occur or later in pregnancy when the placenta takes over hormonal control. CL formation and regression are accompanied by major remodeling of the tissue^18,19,41,42^, which can impact tissue elasticity. Studies suggest that post-ovulatory tissue remodeling is altered with age, evidenced by reduced CL vasculature, alterations in ECM content, and decreased cell proliferation^8,12^. We also observed a decrease in vasculature and a distinct change in the collagen-III to collagen-I ratio, with aged CLs showing almost no collagen III and only a slight reduction in collagen-I. This finding contrasts with reports in the literature, which are mainly based on Sirius Red staining that targets multiple collagen types and was not specifically analyzed for CLs^8,12^. Our findings align with gene expression studies reporting a decrease in collagen-I expression in aged ovaries^37^. Alterations of collagen-I and -III content have been shown to vary with ageing and are dependent on the tissue type^43–45^. Besides overall collagen content, fiber alignment, and topography are crucial factors in determining ECM mechanical properties. Increased collagen fiber thickness and pore size alterations have been linked to tissue stiffening in human ovarian tissues^11^.

It is important to consider the dynamic nature of the tissue. In our study, the heterogeneity in cellular and ECM composition was notable not only between the different age groups but also within one organ. For example, we observed CLs containing microvessels and CLs lacking microvessels side by side in the same ovary. Examining an ovary at a certain age provides only a snapshot in time of an ever-changing organ. Changes in the cellular and molecular composition of CLs are not only linked to ageing but also to the estrous cycle, including CL development and regression^18,19,46^. The variability we observed might be due to the presence of CLs at different stages. Multiple studies have drawn parallels between the processes of ovulation, luteinization and wound healing and scar formation as the CL regresses^47–49^. The complex interplay of ECM composition, fiber alignment, and tissue remodeling during different stages of CL formation and regression, along with the changes that occur during aging, all contribute to tissue elasticity heterogeneity.

We observed minimal changes in follicle elasticity with age or size growth, in contrast with an indentation-based study on bisected organs where higher elasticity was attributed to the presence of larger follicles^23^. However, that study examined larger tissue regions and did not use one-to-one co-registration between tissue features and elasticity, which can likely explain the differences. More prominently, we observed distinct spatial patterns of elasticity in late-stage (larger) follicles. Typically, follicle development is classified in smaller increments from primordial to primary, secondary, pre-antral, and antral stages, with classification in tissue slices usually based on follicle diameter. The QME resolution in this study allowed confident separation only into early or late stages, categorized as ‘small’ or ‘large.’ We selected these categories using an area threshold related to the size of secondary follicles at approximately 200 μm in diameter. At this stage of development, the theca cell layer begins to form surrounding the follicle, with the number of layers increasing as the follicle develops^29^. We observed a stiff shell at the outer edge of the follicles, in line with recent findings using Brillouin microscopy that revealed distinct viscoelastic properties between the outer theca layer and the granulosa cells within the follicle^25^. The stiff shell may be associated with the formation of the theca layer, which consists of an inner vascular-rich theca interna and an outer α-SMA-rich theca externa. Theca cells are highly contractile cells that have been shown to exert compressive stress^50^ onto the follicles, contributing to ovulation^51^. The stiff region coincides with the presence of F-actin and α-SMA, as well as the presence of various ECM molecules such as collagens and fibronectin^50,52^, all of which have been reported to be linked to the cell mechanical phenotype^53–55^. These results suggest that the formation of a stiff theca cell shell may act as a mechanical cage to protect the follicles and oocytes from excessive deformation during development^50^.

## Conclusion

In summary, QME enabled us to visualize and quantify spatial variations in tissue elasticity, revealing non-uniform elasticity across different ovarian compartments. For the first time, we showed that follicles and CLs have distinct elasticity patterns that evolve differently during ageing. By combining QME with immunofluorescence in the same ovary, we were able to link elasticity and volume measurements to the cellular phenotypes and matrix composition of the tissue. Our findings underscore the importance of considering local mechanical environments within the tissue, which may have significant implications for ovarian function and pathology.

### Limitations of the study

High heterogeneity in biological samples, particularly notable in the aged cohort, poses a challenge for drawing conclusive findings with the limited sample size of the current study. Ovarian tissue is highly dynamic, undergoing significant changes during each cycle, resulting in even greater heterogeneity than most other tissues. We propose that specific changes in ECM and cytoskeleton dynamics may contribute to age- and size-dependent elasticity heterogeneities within CLs and follicles during development and aging. Future studies will benefit from a broader selection of markers to comprehensively investigate ECM and cellular changes. Incorporating perturbations such as knockout mouse models or pharmacological inhibitions will further clarify the relationships between cell-molecular phenotypes and tissue elasticity. Additionally, combining QME with complementary microscopy techniques like polarized light microscopy or 3D confocal microscopy of the whole tissue^36^ can enhance our understanding of tissue fiber structure and composition. Although the spatial resolution of current QME measurements was sufficient for outlining ovarian functional structures, higher resolution methods will be essential for more detailed analyses. The perspective of integrating high-resolution variants of QME^56^ with electron microscopy or imaging-based spatial transcriptomics^57^ holds promise for studying cellular and matrix changes in greater detail.

## ACKNOWLEDGMENTS

The Chan lab is supported by the Ministry of Education under the Research Centres of Excellence programme through the Mechanobiology Institute and the Department of Biological Sciences at the National University of Singapore, and the Bia-Echo Asia Centre for Reproductive Longevity & Equality (ACRLE) at the National University of Singapore. C.J.C. also acknowledges the support of the Singaporean Teaching and Academic Research Talent Inauguration Grant (START). A.J. receives a Walter Benjamin fellowship from the Deutsche Forschungsgemeinschaft (DFG, German Research Foundation, Germany; 521343357). B.F.K. acknowledges funding from the Australian Research Council, The Ian Potter Foundation and Department of Health, Western Australia. B.F.K also acknowledges funding from the NAWA Chair programme of the Polish National Agency for Academic Exchange and from the National Science Centre, Poland. We thank Arikta Biswas, Apoorva Shivankar, and Yuting Lou for their discussions and insightful feedback. We thank Hagen Eckert for assistance with the 3D reconstruction.

## AUTHOR CONTRIBUTIONS

Conceptualization, A.J., M.S.H., A.M., C.J.C., B.F.K.; Methodology, M.S.H., A.J., A.M., B.F.K.; Formal Analysis, M.S.H., A.J.; Investigation, A.J, M.S.H.; Writing – Original Draft, A.J., M.S.H.; Writing – Review & Editing, A.J., M.S.H., A.M., B.F.K, C.J.C.; Visualization A.J.; Funding Acquisition, B.F.K., C.J.C.; Resources, B.F.K., C.J.C.; Supervision, B.F.K., C.J.C.

## DECLARATION OF INTERESTS

The authors declare no competing interests.

## Supplementary Figures

**Supplementary Figure S1.**
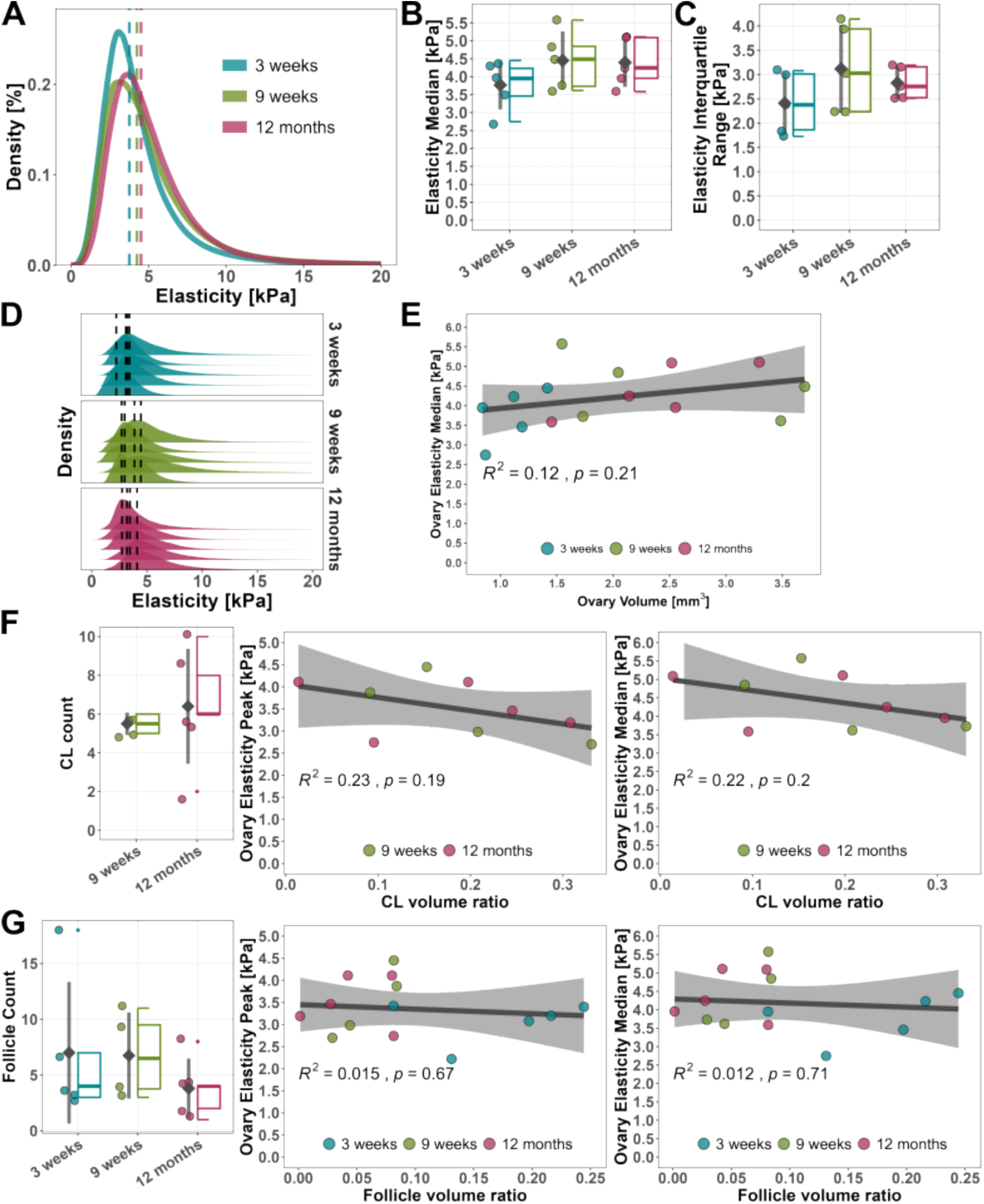
Total murine ovary elasticity at different reproductive stages: A) Distribution of elasticity measurements of all ovaries pooled per age group; dashed lines indicate median elasticity for each age group. B) Elasticity median and C) interquartile range for each ovary. D) Elasticity distributions of each ovary across the different age groups. E) Correlation between median organ elasticity and organ volume within the different age groups. F) CL number (left panel), correlation between peak organ elasticity and CL volume ratio (middle panel), correlation between median organ elasticity and CL volume ratio (right panel) in 9-week and 12-month-old ovaries. G) Follicle number (left panel), correlation between peak organ elasticity and follicle volume ratio (middle panel), correlation between median organ elasticity and follicle volume ratio (right panel) in all age groups. A-G) n = 5 ovaries per age group (3 weeks, 12 months), n = 4 ovaries per age group (9 weeks). B,C,G) Statistical significance between mean values determined by a one-way ANOVA. F) Statistical significance between mean values determined by a Student’s t-test. B,C,F,G) No significant differences were found between the mean values of the age groups.

**Supplementary Figure S2.**
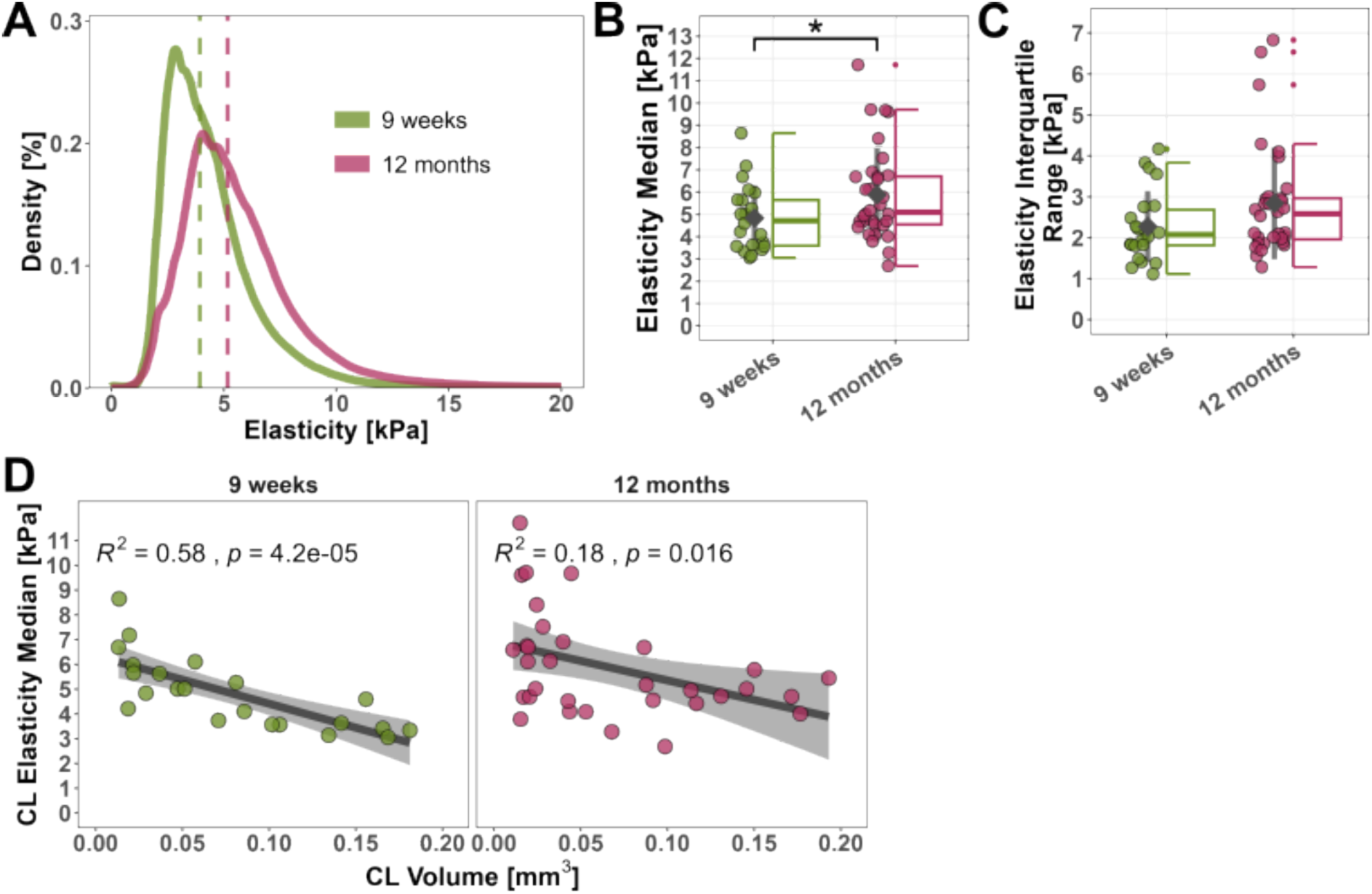
Increased elasticity of CLs in aged ovaries: A) Distribution of elasticity measurements of all CLs pooled per age group; dashed lines indicate median elasticity for each age group. B) Elasticity median and C) interquartile range for each CL. D) Correlation between CL peak elasticity and CL volume. A-D) n = 22 CLs from 4 ovaries (9 weeks) and 32 CLs from 5 ovaries (12 months). B,C) Statistical significance between mean values determined by a Student’s t-test (*p < 0.05). C) No significant differences were found between the mean values of the age groups.

**Supplementary Figure S3.**
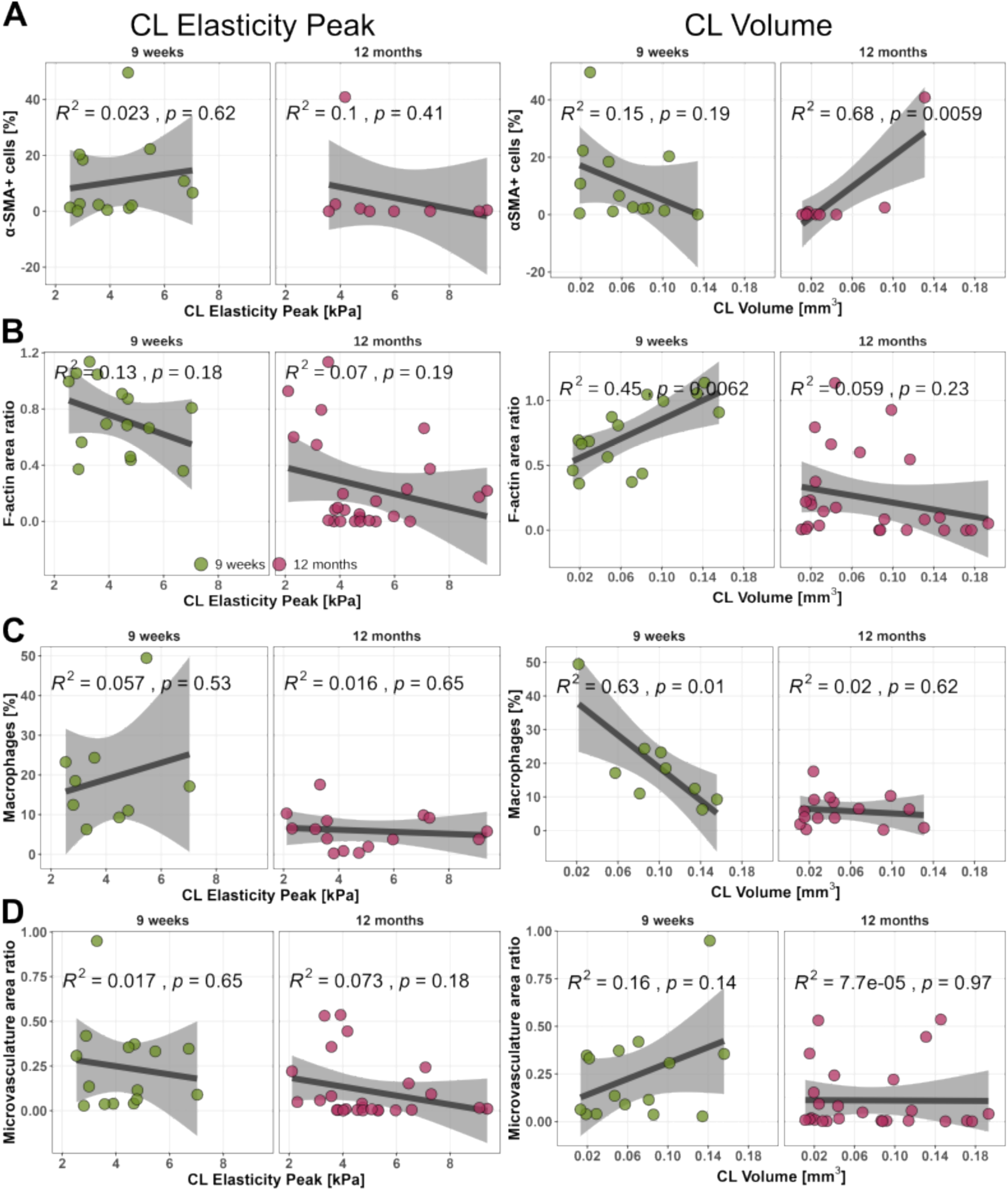
Correlation between cellular phenotypes and CL peak elasticity or CL volume: Trends of changes in A) Percentage of αSMA expressing cells in young and aged CLs, n = 13 CLs from 3 ovaries (9 weeks) and 9 CLs from 2 ovaries (12 months); B) F-actin positive area in young and aged CLs, n = 15 CLs from 3 ovaries (9 weeks) and 26 CLs from 4 ovaries (12 months); C) Percentage of macrophages (F4/80 expressing cells) in young and aged CLs; n = 9 CLs from 2 ovaries (9 weeks) and 15 CLs from 3 ovaries (12 months); D) Microvasculature (CD31) positive area in young and aged CLs, n = 15 CLs from 3 ovaries (9 weeks) and 26 CLs from 4 ovaries (12 months); with elasticity (left panel) and volume (right panel). B,D) Area ratios of markers normalized by the area ratio occupied by nuclei.

**Supplementary Figure S4.**
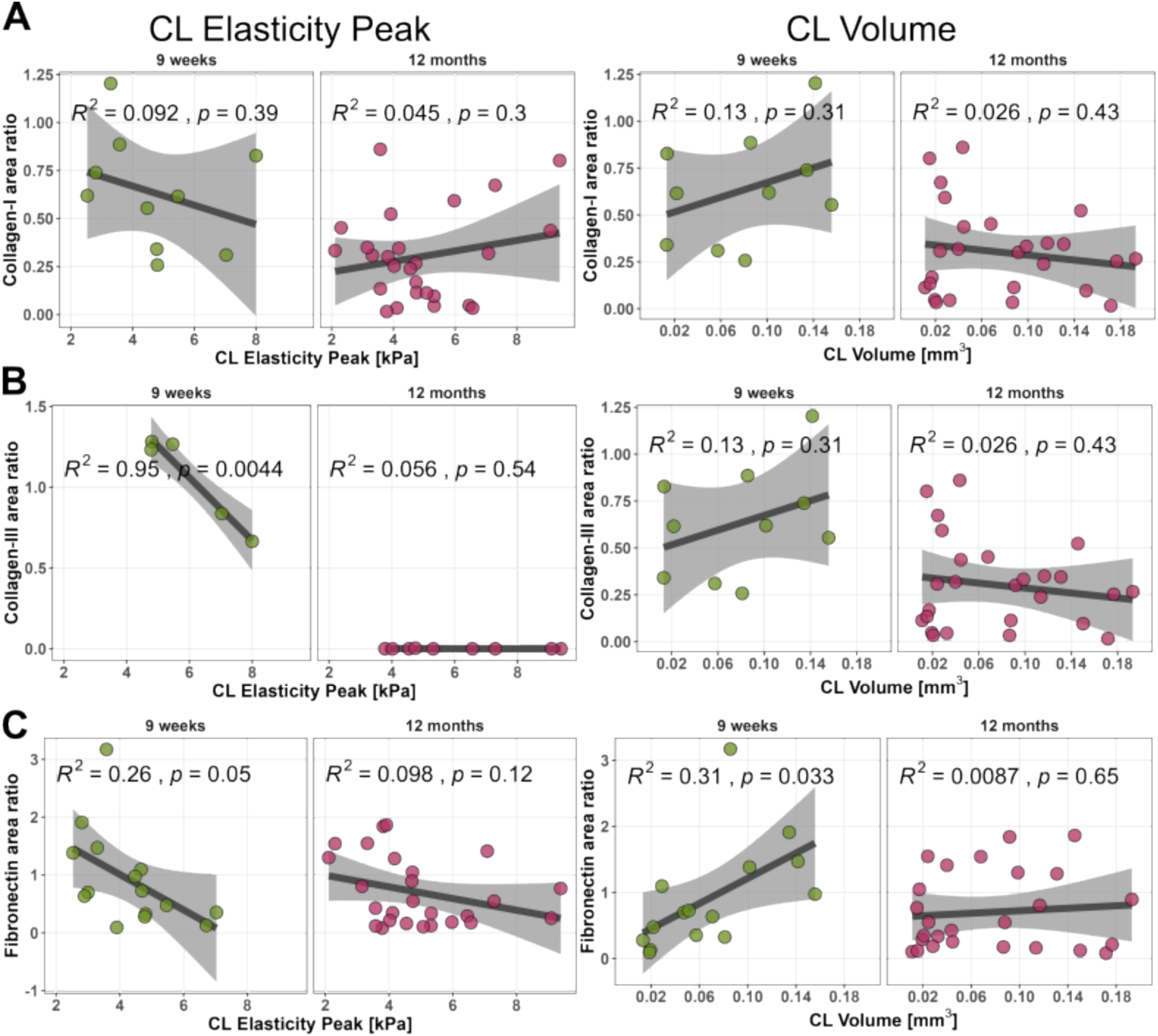
Correlation between matrix composition and CL peak elasticity or CL volume: Trends of changes in A) Collagen-I positive area in young and aged CLs, n = 10 CLs from 2 ovaries (9 weeks) and 26 CLs from 4 ovaries (12 months); B) Collagen-III positive area in young and aged CLs, n = 5 CLs from 1 ovary (9 weeks) and 9 CLs from 2 ovaries (12 months); C) Fibronectin positive area in young and aged CLs, n = 15 CLs from 3 ovaries (9 weeks) and 26 CLs from 4 ovaries (12 months). A-C) area ratios of markers normalized by the area ratio occupied by nuclei.

**Supplementary Video 1:** Video of the QCT and QME image stacks of a 3-week-old murine ovary. https://figshare.com/s/1c762e8df917c1928b45

**Supplementary Video 2:** Video of the QCT and QME image stacks of a 9-week-old murine ovary. https://figshare.com/s/b8487dbd02778cceeca3

**Supplementary Video 3:** Video of the QCT and QME image stacks of a 12-month-old murine ovary. https://figshare.com/s/5161431c6baeb49ebd3c

**Supplementary Video 4:** Animation of the 3D reconstruction of QME data of a 3-week-old murine ovary. https://figshare.com/s/bb1dd2d3c69f8fd6ca8c

**Supplementary Video 5:** Animation of the 3D reconstruction of QME data of a 9-week-old murine ovary. https://figshare.com/s/353d80c176926ff8c6c8

**Supplementary Video 6:** Animation of the 3D reconstruction of QME data of a 9-week-old murine ovary. https://figshare.com/s/a51aed6a4547a0f6dc36

## Methods

## RESOURCE AVAILABILITY

### Lead contact

Further information and requests for resources and reagents should be directed to and will be fulfilled by the Lead Contact, Chii Jou Chan.

## METHOD DETAILS

### Mice

Ovarian tissue was harvested from reproductively young (8 - 10 weeks) and aged (12 – 14 months) female WT IcrTac:ICR, sourced from InVivos (Singapore). All animals were maintained in a specific pathogen-free animal facility at Comparative Medicine (National University of Singapore) and handled according to protocols approved by the institutional animal care and use committees (IACUC; protocol number R22-0511) of the National University of Singapore. Food and water were provided *ad libitum*.

### Ovary encapsulation for QME

Accurate QME measurements require uniaxial compression throughout the sample, which is challenging in small tissues, such as murine ovaries, which feature uneven surfaces. QME uses a bulk pre-strain, *ε*, to ensure uniform contact between the imaging window, compliant layer, sample, and axial translation stage. Small tissues with uneven surfaces require large pre-strain, possibly resulting in QME over-estimating elasticity. By encapsulating the ovaries in agarose hydrogels, we created samples with flat surfaces and controlled thicknesses, enabling uniform contact at low pre-strain and allowing a consistent pre-strain for each sample, which is critical for accurate elasticity estimation and longitudinal comparison. Furthermore, encapsulating the ovaries in a hydrogel also ensures that the tissue remains hydrated throughout the scan acquisition, which is essential for live samples. In this study, the ovaries were encapsulated in 1% (w/v) low-gelling temperature agarose in phosphate-buffered saline (PBS) using 3D-printed (Formlabs) cylindrical plastic molds. The encapsulation resulted in samples with flat surfaces and a controlled height and diameter of 10 mm, essential for accurate QME measurements.

### Quantitative micro-elastography (QME)

#### QME system

Figure 7 shows a schematic of the QME system. The QME measurements were acquired using a fiber-based spectral-domain OCT system (TEL220, Thorlabs) as described previously^38^. The light source was a superluminescent diode with a mean wavelength of 1300 nm and a 3 dB spectral bandwidth of 170 nm. The axial resolution, measured in air, was 4.8 μm (FWHM). The scan lens (LSM03, Thorlabs) used has a numerical aperture of 0.063 and a measured lateral resolution of 7.2 μm (FWHM).

**Figure 7:**
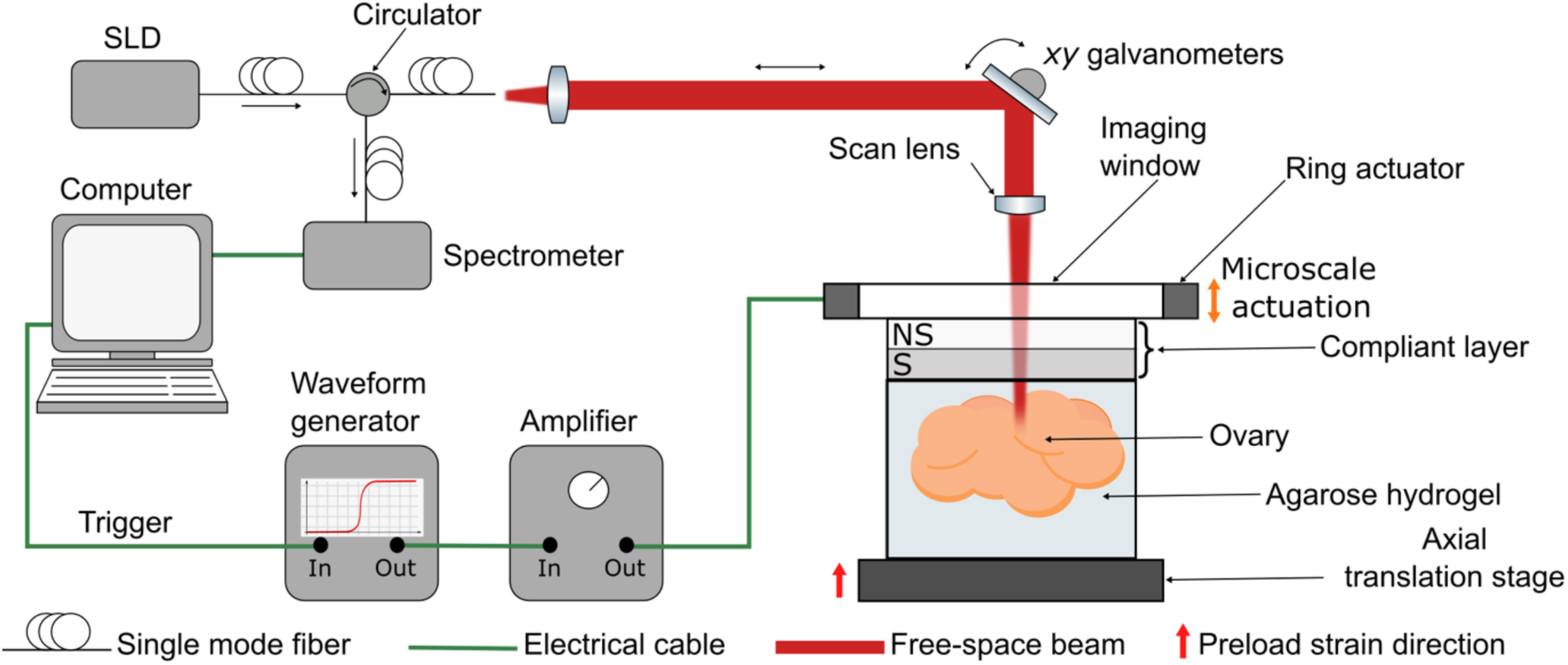
QME experiment setup using a phase-sensitive OCT system in a common-path configuration and compression applied from an annular actuator. SLD: superluminescent diode; NS: non-scattering; S: scattering.

To perform QME, firstly, a compliant silicone layer (P7676 2:1:0.3 crosslinker:catalyst:silicone oil ratio) with a total thickness of approximately 500 μm was placed between the sample surface and imaging window (Edmund Optics) to estimate local axial stress applied to the sample surface^26^. Silicone oil (AK50) was applied to the stress layer-imaging window interface to reduce friction. The compliant layer comprises two sub-layers: a non-scattering layer bonded to a scattering layer, each approximately 250 μm. A pre-strain of 10% of the sample and compliant layer initial height, was applied using a motorized axial translation stage (MLJ050, Thorlabs) to ensure uniform contact between the sample, compliant layer, and imaging window. This resulted in a measured pre-strain of approximately 5% in the compliant layer for each sample. While the sample was under the pre-strain, an additional microscale compression was imparted by applying a stroke of 4 µm using an annular piezoelectric actuator (Piezomechanik GmbH) fixed to a glass imaging window (75 mm diameter). This resulted in local axial strain measurements of approximately -1 to -5 mε in both the compliant layer and tissue. The pre-strain at each lateral location in the compliant layer, was determined by bringing the sample and compliant layer ensemble into contact with the imaging window and acquiring an OCT volume. The initial non-scattering sub-layer thickness, *LI*, was measured by identifying the interface between the non-scattering and scattering layers in each B-scan using an automatic algorithm based on the Canny edge detector in post-processing^58^. A 5% pre-strain of the sample and compliant layer ensemble was applied using the axial translation stage, and another OCT volume was acquired. The final non-scattering sub-layer thickness, *Lf*, was determined using the above-mentioned approach. At each lateral location, *ε* was computed as the change in layer thickness of the final non-scattering layer divided by the initial thickness of the non-scattering layer, i.e., *ε* = *Lf*/*LI* – 1. The local axial strain in the compliant layer resulting from the microscale actuation was measured from the gradient of axial displacement with depth in the scattering region. The local axial stress was then estimated by multiplying the local axial strain in the sub-layer by the tangent modulus (i.e., the gradient of the tangent of the stress-strain curve) at the pre-strain.

The silicone stress-strain relationship was characterized before the QME experiment using a custom-made uniaxial compression testing apparatus described previously^59^. Assuming uniaxial stress, the tangent modulus in the sample is calculated by dividing the local axial strain in the sample by the local axial stress at the sample surface^26^. At pre-strains less than 10%, the tangent modulus is assumed to be equivalent to Young’s modulus.

#### QME acquisition and data processing

QME was performed in a common-path configuration where the interface between the imaging window and compliant layer provides the OCT reference reflection. Each QME scan volume was acquired with 1,000 A-scans per B-scan and 2,000 B-scans over a 4×4 mm (*xy*) lateral region. Two B-scans were acquired for each *y*-location to acquire alternate B-scans at different compression levels. This resulted in a lateral sampling density of 4 μm per voxel in both *x* and *y* The voxel size in *z*, determined by the spectrometer specifications and tissue refractive index, was 2.5 μm in the tissue, assuming a refractive index of 1.3.

The actuator was driven in a quasi-static regime by a 10 Hz square wave, collinearly with the imaging beam and synchronized with the B-scan acquisition. The local axial displacement, *uz*, was calculated from the OCT phase difference between B-scans acquired at the same *y*-location^60^. The dynamic displacement range was extended using a custom phase unwrapping algorithm^61^. Subsequently, gaussian smoothing was applied to the phase difference to alleviate the effects of noise in the measurements by convolving the phase difference with a 3D Gaussian kernel with FWHM of 35×35×8 μm (*xyz*). The local axial strain, *εzz*, was calculated from the gradient of axial displacement using weighted least squares linear regression over a sliding window on the axial displacement of length, Δ*z* of 50 μm, equivalent to 15 pixels in air^62^. Thereby, the weight assigned to each displacement measurement is based on the corresponding OCT SNR at that location. The degradation of axial resolution using least squares linear regression is approximately Δ*z*/√2, resulting in an elasticity system resolution of approximately 35×35×35 μm (*xyz*)^38^. The measured displacement and elasticity sensitivity of the system used in this study has been rigorously characterized previously and are approximately 3.6 nm and 1 kPa, respectively, at an OCT SNR of 35 dB^63^.

Uneven surface topology and mechanical heterogeneity throughout the tissue, despite encapsulation, can result in regions of positive strain in both the tissue and compliant layer as the compressive load is applied across the entire sample surface^64^. By convention, compression is represented by negative strain and the mechanical model in QME assumes only compressive uniaxial stress and strain. Elasticity measurements corresponding to regions of positive strain in the tissue that violate the mechanical model were filtered out to avoid confounding the subsequent analysis and interpretation.

### Tissue fixation and vibratome sectioning

Immediately after the QME scan, the ovaries were incubated in 4% (v/v) paraformaldehyde (PFA)/PBS for 80 minutes. Following fixation, the tissue was kept in a 15% (w/v) sucrose/PBS solution at 4 °C overnight. On the subsequent day, the solution was replaced with PBS containing 30% (w/v) sucrose and 0.02% (w/v) sodium azide. The samples were stored in this buffer at 4°C until vibratome sectioning was performed. For vibratome sectioning, the agarose was carefully trimmed from the QME samples to maintain the tissue architecture. The ovaries were then re-embedded in a 4% (w/v) agarose/PBS solution by placing them on a thin layer of agarose in a mold and covering them with more agarose. After incubating at 37°C for 30 minutes to ensure the two agarose layers merged, the agarose was hardened at 4°C for 15 minutes. The tissue was subsequently sectioned into 60 – 75 μm sections using a Leica vibratome (VT1200S) at a speed of 0.03 mm/s with an amplitude of 2 mm.

### Light microscopy and QME co-registration

For the initial QME co-registration, microscopy images of tissue sections were acquired with a brightfield Echo Revolve Upright Microscope using an Olympus UPlanSApo 10x/0.4 objective. 15 – 25 tiles were captured per section and manually stitched using Fiji (ImageJ v 1.54f)^65^ and the MosaicJ plugin^66^. Structures of interest, follicles, and CLs were manually annotated.

### Segmenting ovarian structures

We used a script written in MATLAB 2023a to extract the volume and elasticity of ovaries, follicles, and CLs. Firstly, the location of the structure of interest, follicle, or CL, was obtained from the annotated bright-field microscopy image. After loading the entire OCT dataset into MATLAB, the corresponding *z*-depth was located in the OCT data set by manually searching for the closest match between the two images. Once the *z*-depth was determined, one x*z* B-scan and one y*z* B-scan were selected from the center of the structure of interest. In each cross-section, a two-dimensional (2D) ROI was manually drawn at the boundary of the structure creating a 2D binary mask in that plane. Each 2D binary mask was extended along the remaining dimension and then multiplied with each other to form a 3D binary mask. The 3D binary mask is then applied to both the OCT SNR and corresponding elasticity data to extract a region corresponding to the structure. The 3D masks used to segment the features were determined from the OCT structural images as the feature boundaries in the OCT images have higher resolution than the corresponding elasticity images. The volume values were computed by multiplying the number of voxels inside the 3D region by the voxel size (4×10^-8^ mm^3^). Once the mask was applied to the elasticity data set, an array of elasticity data (one measurement per voxel) was exported.

Elasticity quantification of the follicle compartments (outer, inner, oocyte) was performed on 2D cross-sections, as volumetric segmentation using concentric spherical masks was highly complex and robustness could not be ensured. A 2D *xy* cross-section was placed through the center of the follicle with the oocyte in focus in the OCT images. Manually drawn regions, approximately circular, were used to segment the follicle compartments. Similar to the 3D segmentation, separate binary masks corresponding to the outer, inner, and oocyte regions were generated and applied to the OCT and elasticity data sets. The area was computed by multiplying the number of pixels inside the 2D region by the pixel size (1.6×10^-5^ mm^2^), and an array of elasticity data was exported.

### Immunofluorescence staining

The tissue sections were stored in PBS containing 0.02% (w/v) sodium azide until staining. For IF staining, sections were transferred to PBS containing 0.5% (v/v) Tween-20 for 5 minutes. For blocking and permeabilization, the samples were incubated in a buffer consisting of 3% BSA (w/v) in PBS with 0.1 M glycine, 0.05% Tween-20, and either 0.3% Triton-X100 (for FN, collagen-I, and collagen-III, α-SMA, F4/80 antibodies) or 0.1% Triton-X100 (for CD31 antibody) for 2 hours at room temperature. Tissue sections were incubated overnight at 4°C in primary antibody solutions, diluted in PBS containing 1% BSA, 0.01 M glycine, 0.05% Tween-20, and 0.02% Triton-X100 (CD31 1:100) or 0.1% Triton-X100 (FN 1:100, collagen-III 1:100, α-SMA 1:500. F4/80 1:50). Following this, the samples were washed four times for 15 minutes each in 1% BSA (w/v) in PBS with 0.05% Tween-20. Secondary antibodies were diluted 1:500 in 1% BSA (w/v) in PBS with 0.05% Tween-20 and staining was performed for 3 hours at room temperature. When collagen-I co-staining was not performed, F-actin was stained using Alexa Fluor 633 or Alexa Fluor 488 conjugated phalloidin (at 1:400 or 1:1000, respectively) alongside 4′,6-diamidino-2-phenylindole (DAPI, nuclear stain; 5 μg/mL) and secondary antibody. After the secondary antibody incubation step, samples were washed three times in 1% BSA/PBS for 15 minutes each, and a final wash was done in PBS for 15 minutes. For collagen-I co-staining, the samples were blocked with rabbit IgG (diluted 1:30 in PBS) for 1 hour at room temperature, followed by overnight incubation with Alexa Fluor 647 conjugated antibody (diluted 1:50 in 0.3% Triton-X100, 1% BSA in PBS) at 4°C. After three washes in 1x PBS for 15 minutes each, DAPI and phalloidin staining were performed for 1 hr at room temperature, followed by 3 more washing steps of 1x PBS for 15 minutes each. All incubation steps were carried out at 50 rpm on a horizontal shaker. Stained sections were mounted on glass microscopy slides using ProLong™ Gold Antifade Mountant.

Confocal microscopy images of fixed samples were acquired on a Zeiss LSM 980 confocal inverted microscope with Plan-Apochromat 20x/0.8 objective. Image stacks were acquired over the entire tissue section with an axial step size of 2 μm and automatic tile stitching with an overlay of 15%.

## QUANTIFICATION AND STATISTICAL ANALYSIS

### Quantification of ECM and cytoskeleton area ratios in CLs

All IF images were processed using Fiji (ImageJ v1.54h)^65^. Firstly, the 3D image stacks were projected into a 2D maximum intensity projection over the whole stack size. A rolling ball algorithm, with a radius of 10 pixels, was applied to correct for an unevenly illuminated background. Noise reduction was achieved using a median filter with a radius of 2 pixels, followed by a Gaussian Blur with a sigma of 0.5 µm. Following the pre-processing, a manual region of interest (ROI) was traced around each CL using the nuclei stain and either the F-actin or collagen stains as informative markers. The ROIs were labeled following the QME co-registration labels, enabling the tracking of the elasticity measurements and the IF data for each ROI. All ROIs were combined for each image, and the information outside the ROIs was excluded before binarization. All ROIs were combined for each image, and the information outside the ROIs was excluded before binarization. A dynamic threshold calculation was applied using the Otsu thresholding algorithm^67^ to convert the greyscale images into binary images. The area ratio for each channel and each ROI was determined using Fiji’s “measure” function. To ensure comparability, the area ratio for each molecule of interest was normalized by the area ratio of the DAPI signal for each ROI.

### Quantification of cell numbers in CLs

Cell counts were obtained using QuPath v0.5.0^68^, where the cell detection function was applied. Separate detection settings were established for each marker based on a representative image, and these same settings were consistently applied to all images. Regions of interest (ROIs) were drawn in Fiji (ImageJ v1.54h)^65^ and directly imported into QuPath for analysis. The number of marker-positive cells was divided by the total number of nuclei to calculate a relative ratio.

### Statistical analysis

All data and statistical analysis were performed in R v4.3.2 and 4.4.0^69^ with the following packages attached: ggplot2^70^, gghalves, ggridges, ggpubr, dplyr, tidyr^71^ or MATLAB v R2023a. The QME elasticity data was described by calculating the mode (peak), FWHM, median, and IQR from all values per age group or per organ, CL, or follicle. All data sets were tested for normal distribution using a Shapiro-Wilk Normality Test^72^. Normal distribution was assumed with p-values ≥ 0.05. For non-normally distributed and ranked data sets (IF staining data), a Mann-Whitney-U was applied to determine statistical significance. Statistical significance in normally distributed data sets (total organ, CL, follicle elasticity; follicle, CL count) was determined using a one-way analysis of variance (ANOVA) for comparisons between 3 groups or a Student’s t-test for comparisons between 2 groups.

### Graphical Representation

Boxplots: Boxes represent the median and 25/75 percentile; whiskers represent the 10/90 percentile. Single points represent the peak or median values. Grey lines indicate linear regression fit with shading of 95% confidence interval. Diamonds and bars represent the mean ± standard deviation of the measurement. Significances are indicated as follows: *p < 0.05, ** p < 0.01, ***p < 0.001. For visualization of the total organ elasticity distributions, the whole organ QME dataset was randomly sampled using the slice_sample function of the dplyr package in R^69^ and 30% of the complete data set was used for the graphical representation. Illustrations for the graphical abstract and Figure 6 were adapted from Servier Medical Art under CC BY 4.0.

### Key resources table

| REAGENT or RESOURCE | SOURCE | IDENTIFIER |
| --- | --- | --- |
| Antibodies and dyes |  |  |
| 4',6-DIAMIDINE-2'-PHENYLINDOLE DIHYDROCHLORIDE (DAPI) | Sigma Aldrich | D8417-5MG |
| Alexa Fluor™ 488 Phalloidin | Invitrogen (Life Technologies holding Pte Ltd.) | A12379 |
| Alexa Fluor™ 633 Phalloidin | Invitrogen (Life Technologies holding Pte Ltd.) | A2228 |
| Donkey anti-Rabbit IgG (H+L) Highly Cross-Adsorbed Secondary Antibody, Alexa Fluor™ 546 | Invitrogen (Life Technologies holding Pte Ltd.) | A10040 |
| Donkey anti-Rat IgG (H+L) Highly Cross-Adsorbed Secondary Antibody, Alexa Fluor™ 488 | Invitrogen (Life Technologies holding Pte Ltd.) | A21208 |
| Mouse F4/80 Antibody [clone # 521204] | R & D systems (SingLabs Technologies Pte Ltd.) | MAB5580 |
| COL1A1 (E8F4L) XP® Rabbit mAb (Alexa Fluor® 647 Conjugate) | Cell Signalling Technologies (Research Instruments Pte Ltd.) | 72827 |
| Anti-Collagen III antibody [EPR17673] | Abcam | ab184993 |
| Anti fibronectin Antibody | Abcam | ab2413 |
| Anti-CD31 antibody [EPR17260-263] | Abcam | ab222783 |
| Rabbit (DA1E) mAb IgG XP® Isotype Control | Cell Signalling Technologies (Research Instruments Pte Ltd.) | 3900S |
| Chemicals, peptides, and recombinant proteins |  |  |
| Paraformaldehyde Solution 4% in 1x PBS | Chem Cruz Santa Cruz Biotechnology (Axil Scientific Pte Ltd.) | SC-281692 |
| Sucrose Ultra-Pure Grade | 1<sup>st</sup> Base (Axil Scientific Pte Ltd.) | Bio-1090-1kg |
| Agarose, low gelling temperature | Sigma Aldrich | A9414-10G |
| UltraPure™ LMP Agarose | Invitrogen (Life Technologies holding Pte Ltd.) | 16520-100 |
| Bovine Serum Albumin | Sigma Aldrich | A9647 |
| Sodium Azide ReagentPlus<sup>R</sup> | Sigma Aldrich | S2002-25G |
| Triton X-100™ Laboratory grade | Sigma Aldrich | X-100-100ML |
| Tween<sup>R</sup> 20 | Sigma Aldrich | P1379-500ML |
| Glycine Ultrapure grade | 1<sup>st</sup> Base (Axil Scientific Pte Ltd.) | Bio-2085-5kg |
| ProLong™ Gold Antifade Mountant | Invitrogen (Life Technologies holding Pte Ltd.) | P36930 |
| ProLong™ Gold Antifade Mountant with DAPI | Invitrogen (Life Technologies holding Pte Ltd.) | P36935 |
| Phosphate Buffered Saline | Gibco (Fisher Scientific Pte Ltd.) | 18912-014 |
| ELASTOSIL® P 7671 A/B | Wacker Chemie AG, Germany | 60039884 |
| AK50 silicone oil | Wacker Chemie AG, Germany | 70702002 |
| Grey Pro Resin | Formlabs, USA | RS-F2-PRGR-01 |
| Experimental models: Organisms/strains |  |  |
| Female mice | Taconics (In Vivos, Singapore) | IcrTac:ICR |
| Software and algorithms |  |  |
| R | R Core Team<sup>69</sup><br>https://www.r-project.org/ | v4.3.2 and 4.4.0 |
| MATLAB® | MathWorks<br>https://ww2.mathworks.cn/en/products/matlab.html | vR2023a |
| Fiji | ImageJ<sup>65</sup><br>https://imagej.net/software/fiji/downloads | ImageJ v1.54f<br>and 1.54h |
| QuPath | https://qupath.github.io/<sup>68</sup> | v0.5.0 |
| SolidWorks, 2019 | Dassault Systèmes, France | SW2019.30 |
| PreForm version 3 | Formlabs, USA | v3.39 |
| Other |  |  |
| Vibratome | Leica | VT 1200S |
| Inverted confocal microscope | Zeiss | LSM 980 |
| Light microscope | Echo | Revolve Upright Microscope |
| Clear Disposable Base Molds, 15mm × 15 mm | Surgipath® | 3803025 |
| MLJ050/M motorized axial translation stage | ThorLabs, Inc., USA | MLJ050/M |
| Form 2 stereolithography 3-D printer | Formlabs, USA | PKG-F2-WSVC-BASIC |
| Form Cure post-processing unit | Formlabs, USA | FH-CU-01 |
| Telesto II spectral domain optical coherence tomography (OCT) system | ThorLabs, Inc., USA | TEL221 |
| OCT scan lens | ThorLabs, Inc., USA | LSM03 |
| Scan head | ThorLabs, Inc., USA | OCTG13 |
| Scan head holder | ThorLabs, Inc., USA | OCT-STAND/M |
| Personal computer | Dell Inc., USA |  |
| Keysight 33500B series waveform generator | Keysight Technologies, Inc., USA | 33500B |
| High voltage amplifier | Piezomechanik GmbH, Germany | LE200/070<br>EBW |
| Custom-made ring actuator with an internal aperture diameter of 65 mm | Piezomechanik GmbH, Germany | N/A |
| Glass imaging window | Edmund Optics Inc., USA | R5001013340-23038 |

## Notes

### Competing Interest Statement

The authors have declared no competing interest.

## REFERENCES

1. Kinnear, H.M., Tomaszewski, C.E., Chang, F.L., Moravek, M.B., Xu, M., Padmanabhan, V., and Shikanov, A. (2020). The ovarian stroma as a new frontier. Reproduction 160, R25–R39. 10.1530/REP-19-0501.

2. Biswas, A., Ng, B.H., Prabhakaran, V.S., and Chan, C.J. (2022). Squeezing the eggs to grow: The mechanobiology of mammalian folliculogenesis. Front. Cell Dev. Biol. 10, 1038107. 10.3389/fcell.2022.1038107.

3. Kawashima, I., and Kawamura, K. (2018). Regulation of follicle growth through hormonal factors and mechanical cues mediated by Hippo signaling pathway. Syst. Biol. Reprod. Med. 64, 3–11. 10.1080/19396368.2017.1411990.

4. West-Farrell, E.R., Xu, M., Gomberg, M.A., Chow, Y.H., Woodruff, T.K., and Shea, L.D. (2009). The Mouse Follicle Microenvironment Regulates Antrum Formation and Steroid Production: Alterations in Gene Expression Profiles1. Biol. Reprod. 80, 432–439. 10.1095/biolreprod.108.071142.

5. West, E.R., Xu, M., Woodruff, T.K., and Shea, L.D. (2007). Physical properties of alginate hydrogels and their effects on in vitro follicle development. Biomaterials 28, 4439–4448. 10.1016/j.biomaterials.2007.07.001.

6. Choi, J.K., Agarwal, P., Huang, H., Zhao, S., and He, X. (2014). The crucial role of mechanical heterogeneity in regulating follicle development and ovulation with engineered ovarian microtissue. Biomaterials 35, 5122–5128. 10.1016/j.biomaterials.2014.03.028.

7. Hornick, J.E., Duncan, F.E., Shea, L.D., and Woodruff, T.K. (2012). Isolated primate primordial follicles require a rigid physical environment to survive and grow in vitro. Hum. Reprod. 27, 1801– 1810. 10.1093/humrep/der468.

8. Amargant, F., Manuel, S.L., Tu, Q., Parkes, W.S., Rivas, F., Zhou, L.T., Rowley, J.E., Villanueva, C.E., Hornick, J.E., Shekhawat, G.S., et al. (2020). Ovarian stiffness increases with age in the mammalian ovary and depends on collagen and hyaluronan matrices. Aging Cell 19, e13259. 10.1111/acel.13259.

9. Briley, S.M., Jasti, S., McCracken, J.M., Hornick, J.E., Fegley, B., Pritchard, M.T., and Duncan, F.E. (2016). Reproductive age-associated fibrosis in the stroma of the mammalian ovary. Reprod. Camb. Engl. 152, 245–260. 10.1530/REP-16-0129.

10. Ouni, E., Bouzin, C., Dolmans, M.M., Marbaix, E., Pyr Dit Ruys, S., Vertommen, D., and Amorim, C.A. (2020). Spatiotemporal changes in mechanical matrisome components of the human ovary from prepuberty to menopause. Hum. Reprod. Oxf. Engl. 35, 1391–1410. 10.1093/humrep/deaa100.

11. Ouni, E., Peaucelle, A., Haas, K.T., Van Kerk, O., Dolmans, M.-M., Tuuri, T., Otala, M., and Amorim, C.A. (2021). A blueprint of the topology and mechanics of the human ovary for next-generation bioengineering and diagnosis. Nat. Commun. 12, 5603. 10.1038/s41467-021-25934-4.

12. Mara, J.N., Zhou, L.T., Larmore, M., Johnson, B., Ayiku, R., Amargant, F., Pritchard, M.T., and Duncan, F.E. (2020). Ovulation and ovarian wound healing are impaired with advanced reproductive age. Aging 12, 9686–9713. 10.18632/aging.103237.

13. Umehara, T., Winstanley, Y.E., Andreas, E., Morimoto, A., Williams, E.J., Smith, K.M., Carroll, J., Febbraio, M.A., Shimada, M., Russell, D.L., et al. (2022). Female reproductive life span is extended by targeted removal of fibrotic collagen from the mouse ovary. Sci. Adv. 8, eabn4564. 10.1126/sciadv.abn4564.

14. Merriman, J.A., Jennings, P.C., McLaughlin, E.A., and Jones, K.T. (2012). Effect of Aging on Superovulation Efficiency, Aneuploidy Rates, and Sister Chromatid Cohesion in Mice Aged Up to 15 Months1. Biol. Reprod. 86. 10.1095/biolreprod.111.095711.

15. Amargant, F., Vieira, C., Pritchard, M.T., and Duncan, F.E. (2024). Systemic low-dose anti-fibrotic treatment attenuates ovarian aging in the mouse. Preprint at bioRxiv, 10.1101/2024.06.21.600035.

16. Wood, C.D., Vijayvergia, M., Miller, F.H., Carroll, T., Fasanati, C., Shea, L.D., Catherine Brinson, L., and Woodruff, T.K. (2015). Multi-modal magnetic resonance elastography for noninvasive assessment of ovarian tissue rigidity in vivo. Acta Biomater. 13, 295–300. 10.1016/j.actbio.2014.11.022.

17. Irving-Rodgers, H., Roger, J., Luck, M., and Rodgers, R. (2006). Extracellular Matrix of the Corpus Luteum. Semin. Reprod. Med. 24, 242–250. 10.1055/s-2006-948553.

18. Irving-Rodgers, H.F., Friden, B.E., Morris, S.E., Mason, H.D., Brannstrom, M., Sekiguchi, K., Sanzen, N., Sorokin, L.M., Sado, Y., Ninomiya, Y., et al. (2006). Extracellular matrix of the human cyclic corpus luteum. Mol. Hum. Reprod. 12, 525–534. 10.1093/molehr/gal060.

19. Irving-Rodgers, H.F., Hummitzsch, K., Murdiyarso, L.S., Bonner, W.M., Sado, Y., Ninomiya, Y., Couchman, J.R., Sorokin, L.M., and Rodgers, R.J. (2010). Dynamics of extracellular matrix in ovarian follicles and corpora lutea of mice. Cell Tissue Res. 339, 613–624. 10.1007/s00441-009-0905-8.

20. Silvester, L.M., and Luck, M.R. (1999). Distribution of extracellular matrix components in the developing ruminant corpus luteum: a wound repair hypothesis for luteinization. Reproduction 116, 187–198. 10.1530/jrf.0.1160187.

21. Jaglan, P., Das, G.K., Kumar, B.V.S., Kumar, R., Khan, F.A., and Meur, S.K. (2010). Cyclical changes in collagen concentration in relation to growth and development of buffalo corpus luteum. Vet. Res. Commun. 34, 511–518. 10.1007/s11259-010-9422-1.

22. Nagamatsu, G., Shimamoto, S., Hamazaki, N., Nishimura, Y., and Hayashi, K. (2019). Mechanical stress accompanied with nuclear rotation is involved in the dormant state of mouse oocytes. Sci. Adv. 5, eaav9960. 10.1126/sciadv.aav9960.

23. Hopkins, T.I.R., Bemmer, V.L., Franks, S., Dunlop, C., Hardy, K., and Dunlop, I.E. (2021). Micromechanical mapping of the intact ovary interior reveals contrasting mechanical roles for follicles and stroma. Biomaterials 277, 121099. 10.1016/j.biomaterials.2021.121099.

24. Henning, N.F.C., and Laronda, M. (2021). The Matrisome Contributes to the Increased Rigidity of the Bovine Ovarian Cortex and Provides a Source of New Bioengineering Tools to Investigate Ovarian Biology. SSRN Electron. J. 10.2139/ssrn.3943652.

25. Chan, C.J., Bevilacqua, C., and Prevedel, R. (2021). Mechanical mapping of mammalian follicle development using Brillouin microscopy. Commun. Biol. 4, 1133. 10.1038/s42003-021-02662-5.

26. Kennedy, K.M., Chin, L., McLaughlin, R.A., Latham, B., Saunders, C.M., Sampson, D.D., and Kennedy, B.F. (2015). Quantitative micro-elastography: imaging of tissue elasticity using compression optical coherence elastography. Sci. Rep. 5, 15538. 10.1038/srep15538.

27. Nelson, J.F., Karelus, K., Felicio, L.S., and Johnson, T.E. (1990). Genetic Influences on the Timing of Puberty in Mice. Biol. Reprod. 42, 649–655. 10.1095/biolreprod42.4.649.

28. Diaz Brinton, R. (2012). Minireview: Translational Animal Models of Human Menopause: Challenges and Emerging Opportunities. Endocrinology 153, 3571–3578. 10.2020071612102345900.

29. Magoffin, D.A. (2005). Ovarian theca cell. Int. J. Biochem. Cell Biol. 37, 1344–1349. 10.1016/j.biocel.2005.01.016.

30. van den Hurk, R., Bevers, M.M., and Beckers, J.F. (1997). In-vivo and in-vitro development of preantral follicles. Theriogenology 47, 73–82. 10.1016/S0093-691X(96)00341-X.

31. Braw-Tal, R., and Yossefi, S. (1997). Studies in vivo and in vitro on the initiation of follicle growth in the bovine ovary. Reproduction 109, 165–171. 10.1530/jrf.0.1090165.

32. Parker, K.J., Doyley, M.M., and Rubens, D.J. (2011). Imaging the elastic properties of tissue: the 20 year perspective. Phys. Med. Biol. 56, 513–513. 10.1088/0031-9155/56/2/513.

33. Mariappan, Y.K., Glaser, K.J., and Ehman, R.L. (2010). Magnetic resonance elastography: A review. Clin. Anat. 23, 497–511. 10.1002/ca.21006.

34. Zaniker, E.J., Zhang, M., Hughes, L., La Follette, L., Atazhanova, T., Trofimchuk, A., Babayev, E., and Duncan, F.E. (2024). Shear wave elastography to assess stiffness of the human ovary and other reproductive tissues across the reproductive lifespan in health and disease. Biol. Reprod., ioae050. 10.1093/biolre/ioae050.

35. Prevedel, R., Diz-Muñoz, A., Ruocco, G., and Antonacci, G. (2019). Brillouin microscopy: an emerging tool for mechanobiology. Nat. Methods 16, 969–977. 10.1038/s41592-019-0543-3.

36. Feng, Y., Cui, P., Lu, X., Hsueh, B., Möller Billig, F., Zarnescu Yanez, L., Tomer, R., Boerboom, D., Carmeliet, P., Deisseroth, K., et al. (2017). CLARITY reveals dynamics of ovarian follicular architecture and vasculature in three-dimensions. Sci. Rep. 7, 44810. 10.1038/srep44810.

37. Lliberos, C., Liew, S.H., Zareie, P., La Gruta, N.L., Mansell, A., and Hutt, K. (2021). Evaluation of inflammation and follicle depletion during ovarian ageing in mice. Sci. Rep. 11, 278. 10.1038/s41598-020-79488-4.

38. Lloyd, E.M., Hepburn, M.S., Li, J., Mowla, A., Hwang, Y., Choi, Y.S., Grounds, M.D., and Kennedy, B.F. (2022). Three-dimensional mechanical characterization of murine skeletal muscle using quantitative micro-elastography. Biomed. Opt. Express 13, 5879. 10.1364/BOE.471062.

39. Humphreys, E.M., Ghione, R., Gosden, R.G., Hobson, B.M., and Wide, L. (1985). Relationship between corpora lutea or fetal number and plasma concentrations of progesterone and testosterone in mice. J. Reprod. Fertil. 75, 7–15. 10.1530/jrf.0.0750007.

40. MacDowell, E.C., and Lord, E.M. (1925). The number of corpora lutea in successive mouse pregnancies. Anat. Rec. 31, 131–141. 10.1002/ar.1090310205.

41. Silvester, L.M., and Luck, M.R. (1999). Distribution of extracellular matrix components in the developing ruminant corpus luteum: a wound repair hypothesis for luteinization. Reproduction 116, 187–198. 10.1530/jrf.0.1160187.

42. Irving-Rodgers, H.F., and Rodgers, R.J. (2005). Extracellular matrix in ovarian follicular development and disease. Cell Tissue Res. 322, 89–98. 10.1007/s00441-005-0042-y.

43. Mays, P.K., Bishop, J.E., and Laurent, G.J. (1988). Age-related changes in the proportion of types I and III collagen. Mech. Ageing Dev. 45, 203–212. 10.1016/0047-6374(88)90002-4.

44. Gao, J., Guo, Z., Zhang, Y., Liu, Y., Xing, F., Wang, J., Luo, X., Kong, Y., and Zhang, G. (2023). Age-related changes in the ratio of Type I/III collagen and fibril diameter in mouse skin. Regen. Biomater. 10, rbac110. 10.1093/rb/rbac110.

45. Horn, M.A., and Trafford, A.W. (2016). Aging and the cardiac collagen matrix: Novel mediators of fibrotic remodelling. J. Mol. Cell. Cardiol. 93, 175–185. 10.1016/j.yjmcc.2015.11.005.

46. Iwahashi, M., Muragaki, Y., Ooshima, A., and Umesaki, N. (2006). Immunohistochemical analysis of collagen expression in human corpora lutea during the menstrual cycle and early pregnancy. Fertil. Steril. 85, 1093–1096. 10.1016/j.fertnstert.2005.10.022.

47. Smith, M.F., McIntush, E.W., and Smith, G.W. (1994). Mechanisms associated with corpus luteum development. J. Anim. Sci. 72, 1857–1872. 10.2527/1994.7271857x.

48. Duffy, D.M., Ko, C., Jo, M., Brannstrom, M., and Curry, T.E., Jr. (2019). Ovulation: Parallels With Inflammatory Processes. Endocr. Rev. 40, 369–416. 10.1210/er.2018-00075.

49. Lu, E., Li, C., Wang, J., and Zhang, C. (2019). Inflammation and angiogenesis in the corpus luteum. J. Obstet. Gynaecol. Res. 45, 1967–1974. 10.1111/jog.14076.

50. Biswas, A., Lou, Y., Ng, B.H., Tomida, K., Darpe, S., Wu, Z., Lu, T.B., Bonne, I., and Chan, C.J. (2024). Theca cell mechanics and tissue pressure regulate mammalian ovarian folliculogenesis. Preprint at bioRxiv, 10.1101/2024.05.06.592641.

51. Martin, G.G., and Talbot, P. (1981). The role of follicular smooth muscle cells in hamster ovulation. J. Exp. Zool. 216, 469–482. 10.1002/jez.1402160316.

52. Berkholtz, C.B., Lai, B.E., Woodruff, T.K., and Shea, L.D. (2006). Distribution of extracellular matrix proteins type I collagen, type IV collagen, fibronectin, and laminin in mouse folliculogenesis. Histochem. Cell Biol. 126, 583–592. 10.1007/s00418-006-0194-1.

53. Gavara, N., and Chadwick, R.S. (2016). Relationship between cell stiffness and stress fiber amount, assessed by simultaneous atomic force microscopy and live-cell fluorescence imaging. Biomech. Model. Mechanobiol. 15, 511–523. 10.1007/s10237-015-0706-9.

54. Kaukonen, R., Mai, A., Georgiadou, M., Saari, M., De Franceschi, N., Betz, T., Sihto, H., Ventelä, S., Elo, L., Jokitalo, E., et al. (2016). Normal stroma suppresses cancer cell proliferation via mechanosensitive regulation of JMJD1a-mediated transcription. Nat. Commun. 7, 12237. 10.1038/ncomms12237.

55. Jaeschke, A., Jacobi, A., Lawrence, M.G., Risbridger, G.P., Frydenberg, M., Williams, E.D., Vela, I., Hutmacher, D.W., Bray, L.J., and Taubenberger, A. (2020). Cancer-associated fibroblasts of the prostate promote a compliant and more invasive phenotype in benign prostate epithelial cells. Mater. Today Bio, 100073. 10.1016/j.mtbio.2020.100073.

56. Mowla, A., Li, J., Hepburn, M.S., Maher, S., Chin, L., Yeoh, G.C., Choi, Y.S., and Kennedy, B.F. (2022). Subcellular mechano-microscopy: high resolution three-dimensional elasticity mapping using optical coherence microscopy. Opt. Lett. 47, 3303. 10.1364/OL.451681.

57. He, S., Bhatt, R., Brown, C., Brown, E.A., Buhr, D.L., Chantranuvatana, K., Danaher, P., Dunaway, D., Garrison, R.G., Geiss, G., et al. (2022). High-plex imaging of RNA and proteins at subcellular resolution in fixed tissue by spatial molecular imaging. Nat. Biotechnol. 40, 1794–1806. 10.1038/s41587-022-01483-z.

58. Kennedy, K.M., Es’haghian, S., Chin, L., McLaughlin, R.A., Sampson, D.D., and Kennedy, B.F. (2014). Optical palpation: optical coherence tomography-based tactile imaging using a compliant sensor. Opt. Lett. 39, 3014–3017. 10.1364/OL.39.003014.

59. Sanderson, R.W., Curatolo, A., Wijesinghe, P., Chin, L., and Kennedy, B.F. (2019). Finger-mounted quantitative micro-elastography. Biomed. Opt. Express 10, 1760–1773. 10.1364/BOE.10.001760.

60. Wang, R.K., Ma, Z., and Kirkpatrick, S.J. (2006). Tissue Doppler optical coherence elastography for real time strain rate and strain mapping of soft tissue. Appl. Phys. Lett. 89, 144103. 10.1063/1.2357854.

61. Kennedy, B.F., McLaughlin, R.A., Kennedy, K.M., Chin, L., Curatolo, A., Tien, A., Latham, B., Saunders, C.M., and Sampson, D.D. (2014). Optical coherence micro-elastography: mechanical-contrast imaging of tissue microstructure. Biomed. Opt. Express 5, 2113–2124. 10.1364/BOE.5.002113.

62. Kennedy, B.F., Koh, S.H., McLaughlin, R.A., Kennedy, K.M., Munro, P.R.T., and Sampson, D.D. (2012). Strain estimation in phase-sensitive optical coherence elastography. Biomed. Opt. Express 3, 1865–1879. 10.1364/BOE.3.001865.

63. Li, J., Hepburn, M.S., Chin, L., Mowla, A., and Kennedy, B.F. (2021). Analysis of sensitivity in quantitative micro-elastography. Biomed. Opt. Express 12, 1725–1745. 10.1364/BOE.417829.

64. Wijesinghe, P., Chin, L., and Kennedy, B.F. (2018). Strain tensor imaging in compression optical coherence elastography. IEEE J. Sel. Top. Quantum Electron., 1–1. 10.1109/JSTQE.2018.2871596.

65. Schindelin, J., Arganda-Carreras, I., Frise, E., Kaynig, V., Longair, M., Pietzsch, T., Preibisch, S., Rueden, C., Saalfeld, S., Schmid, B., et al. (2012). Fiji: an open-source platform for biological-image analysis. Nat. Methods 9, 676–682. 10.1038/nmeth.2019.

66. Thévenaz, P., and Unser, M. (2007). User-friendly semiautomated assembly of accurate image mosaics in microscopy. Microsc. Res. Tech. 70, 135–146. 10.1002/jemt.20393.

67. Otsu, N. (1979). A Threshold Selection Method from Gray-Level Histograms. IEEE Trans. Syst. Man Cybern. 9, 62–66. 10.1109/tsmc.1979.4310076.

68. Bankhead, P., Loughrey, M.B., Fernández, J.A., Dombrowski, Y., McArt, D.G., Dunne, P.D., McQuaid, S., Gray, R.T., Murray, L.J., Coleman, H.G., et al. (2017). QuPath: Open source software for digital pathology image analysis. Sci. Rep. 7, 16878. 10.1038/s41598-017-17204-5.

69. R Core Team (2013). R: A Language and Environment for Statistical Computing.

70. Wickham, H. (2016). ggplot2: Elegant graphics for data analysis (Springer-Verlag New York).

71. Wickham, H., Averick, M., Bryan, J., Chang, W., McGowan, L.D., François, R., Grolemund, G., Hayes, A., Henry, L., Hester, J., et al. (2019). Welcome to the tidyverse. J. Open Source Softw. 4, 1686. 10.21105/joss.01686.

72. Royston, J.P. (1982). Algorithm AS 181: The W Test for Normality. J. R. Stat. Soc. Ser. C Appl. Stat. 31, 176–180. 10.2307/2347986.

